# Rab11b-mediated integrin recycling promotes brain metastatic adaptation and outgrowth

**DOI:** 10.1101/666750

**Authors:** Erin N. Howe, Miranda D. Burnette, Melanie E. Justice, James W. Clancy, Ian H. Guldner, Patricia M. Schnepp, Victoria Hendrick, Uma K. Aryal, Alicia T. Specht, Jun Li, Crislyn D’Souza-Schorey, Jeremiah Z. Zartman, Siyuan Zhang

## Abstract

**GRAPHICAL ABSTRACT:** 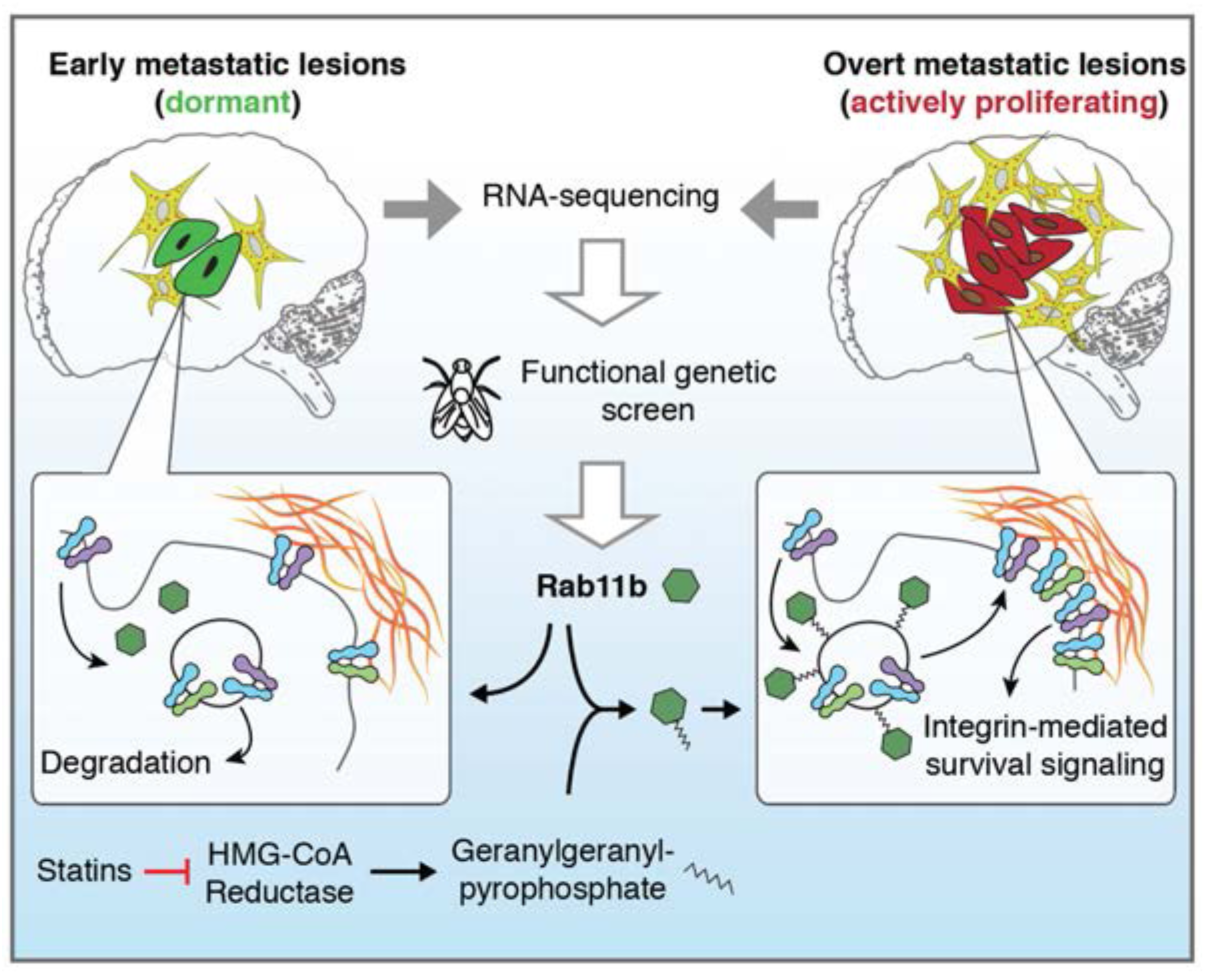

**SUMMARY:** Breast cancer brain metastases (BCBM) have a 5-20 year latency and account for up to 30% of mortality. Developing new therapeutics requires a molecular understanding of adaptation to the brain microenvironment. Here, we combined RNA-sequencing of BCBM development with a reverse genetic screen in Drosophila melanogaster and identified Rab11b, an endosomal recycling protein, as a mediator of metastatic adaptation. We show that disseminated cells up-regulate Rab11b early after arrival in the brain, allowing control of the cell surface proteome through recycling of proteins required for successful interaction with the microenvironment, including integrin β1. Rab11b-mediated control of integrin β1 surface expression allows ligation to the brain ECM, activating mechanotransduction signaling to allow survival and proliferation. We propose a model in which up-regulation of Rab11b allows disseminated cells to recycle needed proteins during metastatic adaptation, without strictly requiring transcription and translation, to allow for metastatic outgrowth.

**Manuscript Summary:** Rab11b up-regulation in the brain microenvironment promotes recycling of cargo proteins required for breast cancer brain metastasis, including increased surface expression of integrin β1, which allows brain extracellular matrix attachment and mechanotransduction. Inhibition of the mevalonate pathway with statins prevents geranylgeranylation of Rab11b, decreasing cargo recycling, and inhibiting brain metastasis.

## INTRODUCTION

Breast cancer brain metastases (BCBM) are an increasingly urgent clinical problem, with patient survival measured in months (Eichler et al., 2011; Frisk et al., 2017; Nam et al., 2008; Niwinska et al., 2017). Systemic treatments, such as chemotherapies or targeted therapies, cannot effectively treat micrometastatic brain lesions or prevent brain metastatic relapse, largely due to their inability to penetrate the blood-brain barrier (BBB) (Carson et al., 2006; Peereboom, 2005). Therefore, with better control of systemic disease, many women who have a stable primary disease, or respond to initial treatment, ultimately develop BCBM. As a result, the BCBM incidence is increasing (Kodack et al., 2015; Kotecki et al., 2018).

Metastasis is an inefficient process, with disseminated tumor cells (DTCs) dying at every stage of the process. Prior to the formation of overt metastatic disease, DTCs persist in the brain for months or years, and effective engagement with the metastatic niche is essential for colonization and outgrowth (Harper et al., 2016; Hosseini et al., 2016; Massagué and Obenauf, 2016; Sosa et al., 2014). While the classic ‘seed and soil’ hypothesis highlights the importance of optimal DTCs arriving in a permissive metastatic microenvironment, recent studies into metastatic seeding, dormany, and outgrowth have revealed dynamic co-evolutionary processes between DTCs and the metastatic niche (Celià-Terrassa and Kang, 2018). Indeed, successful metastatic adaptation requires organ-specific interactions with the surrounding parenchymal cells and extracellular matrix (Obenauf and Massagué, 2015), and the diversity of these interactions (Chen et al., 2016; Contreras-Zárate et al., 2019; Er et al., 2018; Schnepp et al., 2017; Zhang et al., 2015), combined with the evolving heterogeneity of the DTCs themselves, renders the mechanistic dissection required for the development of treatment options challenging. Once DTCs have adapted to the metastatic microenvironment and begun proliferating, current treatments often fail (Achrol et al., 2019); thus, identifying common mechanisms underlying the ability of DTCs to adapt to the metastatic microenvironment is critically important.

The ability of a DTC to successfully engage the metastatic microenvironment is dictated by the composition of the cell surface, which governs ligation of adhesion complexes, binding of growth factors, and engagement with parenchymal cells. Although much work has focused on transcriptional changes during tumorigenesis and metastasis, control of the cell surface through vesicular trafficking is emerging as a mechanism of regulating several hallmarks of cancer (Mosesson et al., 2008). Indeed, vesicular trafficking, including endocytosis and endosomal recycling, is the primary mechanism regulating the composition and organization of the cell surface (Le Roy and Wrana, 2005; Welz et al., 2014). Trafficking controls the localization and function of a variety of surface proteins with known roles in cancer and metastasis, including E-cadherin, EGFR, and integrins (Caswell et al., 2008; Le et al., 1999; Ye et al., 2016). Yet, the role of trafficking, and the central machinery by which DTCs control the surface proteome in response to the metastatic microenvironment remain poorly defined.

In this study, we identify Rab11b-mediated endosomal recycling as a unique mechanism for cancer cell adaptation to a challenging brain metastatic microenvironment. We first identify differentially regulated genes by utilizing RNA-sequencing to identify temporal changes during BCBM development. We then screen those genes for a functional role in brain metastasis using a **Drosophila melanogaster** tumor model (Willecke et al., 2011; Wu et al., 2010), leading to the identification of Rab11b, a mediator of endosomal recycling (Kelly et al., 2012). The Rab11 family of small GTPases is critical for recycling of a number of proteins, and has been implicated in several types of cancer (Chung et al., 2014; Palmieri et al., 2006a; Yoon et al., 2005; Yu et al., 2016). Perhaps the least well-studied family member, Rab11b localizes to the endosomal recycling center (ERC) (Schlierf et al., 2000), and is mainly expressed in non-epithelial tissues, including brain (Lai et al., 1994). We find that breast cancer cells up-regulate Rab11b during early adaptation to the brain metastatic site, providing a mechanism for DTCs to recycle needed proteins during this critical step of the metastatic cascade, enabling survival and outgrowth. Mechanistically, Rab11b-mediated control of the cell surface proteome, including recycling of integrin β1 enables successful interaction with the brain extracellular matrix and mechanotransduction-activated survival signaling. Our findings suggest recycling controls the composition of the cell surface proteome, which is critically important for metastatic cell-microenvironmental interaction and eventual outgrowth.

## RESULTS

### Identification of functional mediators of brain metastasis outgrowth

To dissect temporal changes during breast cancer brain metastatic outgrowth, we analyzed the transcriptional changes between early colonized cells at brain (7 days post injection, dpi) and late-stage overt brain metastases (40 dpi) using RNA-sequencing (Figure 1A). Histology and cytokeratin 8 (K8) staining revealed the relative size of brain metastases and confirm the presence of colonized tumor cells in 7 dpi samples (Figure 1B, black arrows). tdTomato-positive brain metastases were dissected from fresh brain tissue and sequenced. Nine 40 dpi samples were split into three groups based on the size of the brain metastasis at the time of dissection (small, medium, large). To exclude brain tissue-derived reads (mouse origin), only sequencing reads that uniquely mapped to human genome were kept for downstream analysis (Figure S1A, B). We found that the 40 dpi brain metastases clustered away from 7 dpi samples, regardless of the size at dissection (Figure S1C), suggesting that metastatic adaptation and acquisition of a proliferative phenotype directs transcriptional reprogramming. Due to their similarity, we grouped 40 dpi samples together irrespective of size using Fisher’s combined test. We identified 125 genes that were significantly differentially regulated during breast cancer adaptation to the brain metastatic site with a Fisher’s combined p-value < 0.05 (Figure 1C). Of the 125 dysregulated genes, 108 genes up-regulated in overt brain metastasis (40 dpi) (Figure 1D, BrainMets Sig.Genes).

**Figure 1.**
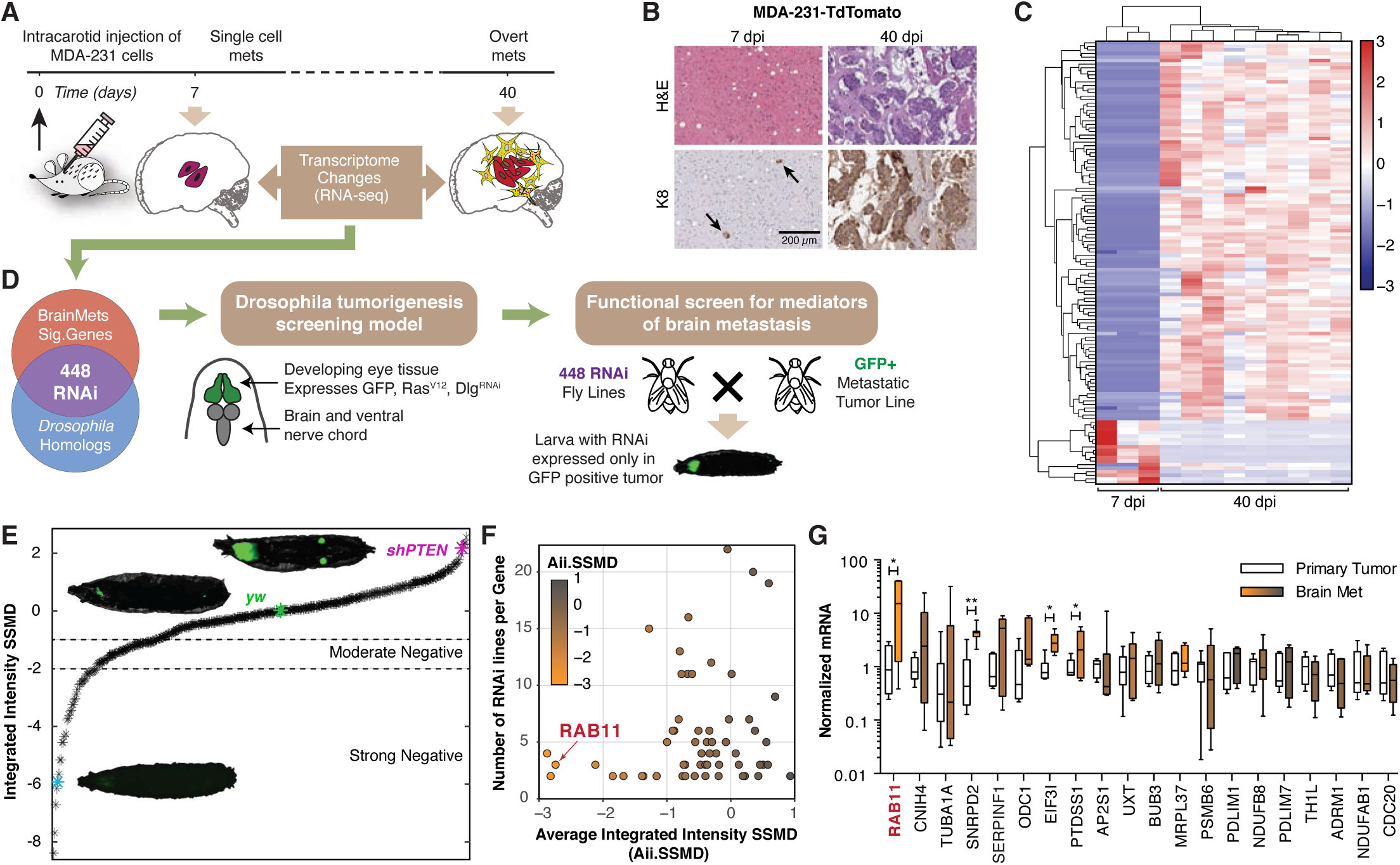
Identification of functional mediators of brain metastasis. **(A)** Schematic of experimental design to generate temporal transcriptome of breast cancer brain metastases. **(B)** Representative images of brain mets at 7 or 40 dpi, as indicated. Tissues were H&E stained and cytokeratin 8 (K8) stained to show tumor tissue. Scale bar is 200 µm. **(C)** Heatmap showing expression of 125 genes differentially expressed between 7 and 40 dpi. Fisher’s combined test, Bonferroni p-value < 0.05. **(D)** Schematic of selection of *Drosophila* homologs, the genotype and phenotype of the *Drosophila* screening line, and the functional screening model used. **(E)** Data is presented as strictly standardized mean difference (SSMD) calculated with respect to the negative control (yw, no RNAi construct) shown in green, and the positive control (shPTEN) shown in purple for a minimum of 15 larvae per cross. Hits are characterized as moderate or strong positives as indicated. Representative images for both controls, and a strong positive hit are shown. **(F)** For each RNAi line that yielded a strong, moderate or weak negative phenotype, the number of RNAi lines for that gene is plotted against the average integrated intensity SSMD. Datapoints are colored by average integrated intensity SSMD, with orange indicating a negative average SSMD, and gray indicating a positive average SSMD. **(G)** qRT-PCR for 20 genes with lowest average integrated intensity SSMD scores in MDA-231 primary mammary fat pad tumors (Primary Tumor, white), and brain metastases (Brain Met, orange to gray). Brain metastasis samples are colored to correspond to the gene’s average integrated intensity SSMD, as in F. *Boxes*, first to third interquartile range, *line*, mean, *whiskers*, minimum and maximum values. Pairwise comparisons made as indicated using Student’s t-test. * p < 0.05, ** p < 0.01.

To identify genes that functionally drive brain metastasis progression, we employed a Drosophila melanogaster tumor model for in vivo screening of the BrainMets Sig.Genes (Figure 1D) (Pagliarini and Xu, 2003; Willecke et al., 2011). This model overexpresses oncogenic Ras^V12^, an RNAi construct targeting the polarity gene discs large (Dlg), and green fluorescent protein (GFP) in the epithelial Drosophila imaginal eye disc. In this model, tumors develop in the eye disc and progressively invade into adjacent brain tissue (Figure 1D, Figure S2B). We identified Drosophila orthologs for the 108 BrainMets Sig.Genes (Hu et al., 2011), and obtained 448 RNAi fly lines (Figure S2A). Male RNAi flies were crossed with virgin female flies from the GFP, Ras^V12^, Dlg^RNAi^ tumor tester line, larvae were collected 6 days post-egg laying, and whole larva brightfield and GFP imaging were performed. For a minimum of fifteen animals per cross, the integrated intensity of the GFP signal within the larval body was measured, and the effect size for each RNAi line was calculated as compared to the negative control RNAi line (yw, green) and the positive control (shPTEN, purple) using the strictly standardized mean difference (SSMD) of integrated fluorescence intensity (Figure 1E). We identified 39 Drosophila RNAi lines representing 29 human genes that strongly suppressed tumor growth (Figure 1E, Strong Negative SSMD, Figure S2C, 962 – Rab11), while an additional 53 RNAi lines presenting 32 human genes moderately suppressed tumor growth (Figure 1E, Moderate Negative SSMD, Figure S2C, 561 – PSMC6). Interestingly, for the BrainMets Sig.Genes screened, gene expression was not strongly correlated with suppression of tumor growth (Figure S2D), suggesting that gene expression alone does not fully predict the importance of a gene in tumorigenesis and metastasis.

To account for the biological variations among RNAi constructs, we averaged SSMD scores for all RNAi lines targeting each gene to identify the genes that consistently decreased tumor growth and metastasis (Figure 1F and Figure S2E). Among the genes that strongly inhibited tumor growth - genes with the lowest average combined integrated intensity SSMD scores (Aii.SSMD) - we found, SERPINF1, a member of the SERPIN family. SERPIN family genes were previously implicated in brain metastasis initiation (Valiente et al., 2014), demonstrating the predictive power of our screening model for metastatic progression (Figure 1F). We identified the top 20 genes based on Aii.SSMD, and examined their expression in samples from MDA-231 human breast cancer xenograft primary and brain metastases. Among the top 3 genes with the lowest Aii.SSMD (RAB11, SNRPD2, MRPL37), RAB11 mRNA increased 18-fold in brain metastasis tissue (Figure 1G). In contrast with SERPINF1, which is moderately up-regulated in brain metastases, the highly inducible nature of RAB11 in the brain metastases suggests a brain context-specific function of RAB11(Figure 1G).

### Rab11b is up-regulated is required for metastatic adaptation to the brain microenvironment

Rab11 is a family of small GTPases that regulate vesicular transport in the endosomal and exosomal recycling pathways, where they recycle numerous important proteins, including E-cadherin, epidermal growth factor receptor (EGFR), fibroblast growth factor receptor (FGFR), and multiple integrins (Caswell et al., 2008; Haugsten et al., 2014; Le et al., 1999; Paterson et al., 2003). Examination of a human tissue microarray containing 39 primary and 10 breast cancer brain metastasis samples confirms higher expression of Rab11 in brain metastases (Figure 2A). While the **Drosophila)** genome encodes a single Rab11 (Sasamura et al., 1997), there are three mammalian Rab11 family members: Rab11a, Rab25 (Rab11c), and Rab11b. Compared to cell line or primary tumors, the mRNA level of Rab11b, but not Rab11a or Rab25, is significantly up-regulated in brain metastases (Figure 2B). Further, brain metastases derived from multiple breast cancer cell lines showed more than 15 fold increase of Rab11b expression compared with cultured cells (Figure 2C, D), with no change in Rab11a or Rab25 levels (Figure S3A), suggesting the brain metastatic microenvironment specifically influences Rab11b expression. To determine if we can recapitulate the induction of Rab11b **in vitro**, we co-cultured breast cancer cells with primary murine glia (Zhang et al., 2015), and found that only Rab11b is up-regulated (Figure 2E, Figure S3B). Co-culture with murine caveolin 1-/- fibroblasts, a model of cancer-associated fibroblast (CAFS) (Sotgia et al., 2009), showed no increase in Rab11b (Figure 2E), confirming that the brain metastatic microenvironment uniquely induces expression of Rab11b.

**Figure 2.**
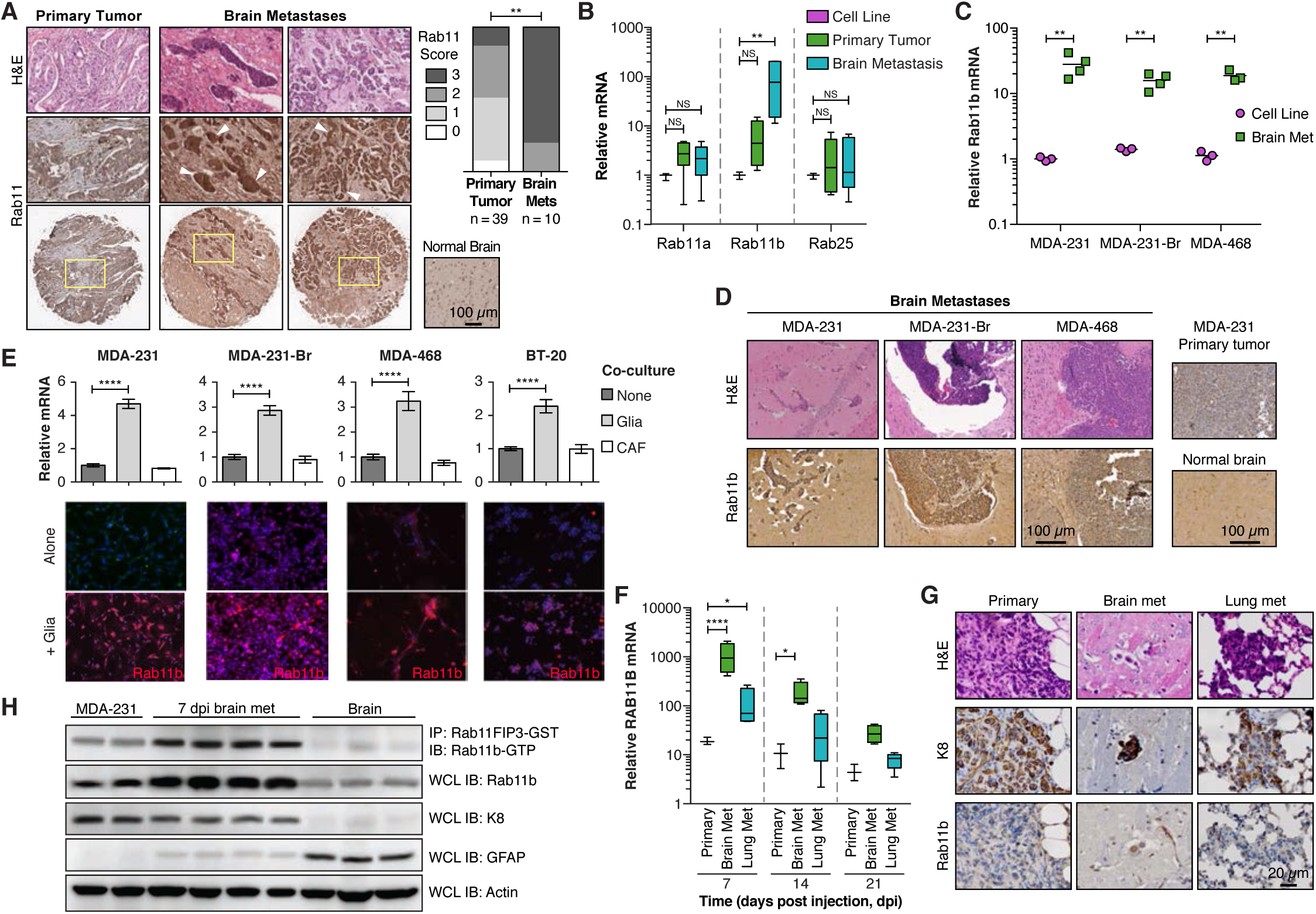
Rab11b is up-regulated during metastatic adaptation to the brain microenvironment. **(A)** Representative images of H&E (top) or Rab11 (middle and bottom) immunohistochemical staining of human primary breast cancer or breast cancer brain metastases (arrowheads). Left, scoring of Rab11 IHC. Analysis of contingency, Fisher’s exact test. **(B)** qPCR for Rab11 isoforms in MDA-231 cells grown in culture, primary tumors (21 dpi), or brain metastases (21 dpi). Values for each isoform are normalized to cells in culture. ANOVA with Dunnett’s multiple comparison test. **(C)** qPCR for Rab11b in cells grown in culture versus brain metastases. All values are normalized to MDA-231 cells in culture. Student’s t-test. **(D)** Representative H&E and Rab11b immunohistochemical staining of brain metastases, and MDA-231 primary tumor or normal murine brain. **(E)** *Top*, qPCR for Rab11b in cell lines co-cultured with primary murine glia or CAF cells for 2 days. All values normalized to single culture. ANOVA with Dunnett’s multiple comparison test. *Bottom*, immunostaining for Rab11b (green, red) and nuclei (DAPI, blue) in cell lines cultured alone or co-cultured with primary murine glia for 5 days. **(F)** qPCR for Rab11b in MDA-231 primary tumors or metastases as indicated, collected at time points indicated. All values are normalized to MDA-231 cells in culture. Two-way ANOVA with Tukey’s multiple comparison test. **(G)** Representative H&E, cytokeratin 8 (K8), and Rab11b immunohistochemical staining of MDA-231 primary tumors, brain or lung metastases, at 7 dpi. **(H)** MDA-231 cells were intracranially injected and brain metastases were dissected at 7dpi using fluorescence signal as a guide. Metastases were dissociated and the bulk of brain parenchymal cells were removed using magnetic bead-based stromal cell depletion. Naive brain (brain), or stroma depleted brain met (7 dpi brain met) samples were lysed and subjected to Rab11b activation assay, followed by immunoblotting. For all panels, * p < 0.05, ** p < 0.01, *** p < 0.001, **** p < 0.0001.

To explore the temporal regulation of Rab11b during tumorigenesis or metastasis, we allowed MDA-231 cells to form primary tumors, lung metastases or brain metastases. We found that Rab11b is up-regulated in all three tissue microenvironments early in tumor or metastasis formation; however, Rab11b is significantly more strongly up-regulated during brain metastasis formation during the early metastatic adaptation stage (Figure 2F, 7 dpi). Immunohistochemistry (IHC) staining of Rab11b in primary tumors and lung or brain metastases derived from MDA-231 confirms high expression of Rab11b protein specifically early during brain metastasis formation (Figure 2G). To confirm that the increased level of Rab11b leads to increased activation of Rab11b, we incubated naïve brain or dissected MDA-231 brain metastasis lysates with Rab11FIP3-GST, which specifically binds Rab11b-GTP (Junutula et al., 2004). We found significantly increased expression of total Rab11b, confirming our IHC results, as well as a strong induction of active Rab11b-GTP (Figure 2H), suggesting that not only is Rab11b expression increased in breast cancer brain metastases, but this increase translates to more active Rab11b.

To determine whether Rab11b plays a causal role in breast cancer brain metastasis formation, we used two shRNA constructs targeting Rab11b and confirmed that they decrease Rab11b mRNA and protein (Figure 3A, B, Figure S4A, B), as well as active Rab11b (Figure 3C). To confirm that shRab11b constructs are able to block glial-mediated up-regulation of Rab11b, cells were co-cultured with primary murine glia, and glia were removed using MACS prior to analysis. Although glia are able to induce up-regulation of Rab11b, the effect is severely blunted in cells expressing shRab11b constructs (Figure 3D, E, Figure S4C, D). Depletion of Rab11b dramatically decreases brain metastasis in both intracranial (Figure 3F, Figure S4E), and intracardiac (Figure 3G, Figure S4F) injection models, with a significant reduction in incidence and proliferative ability (Figure 3H,I). Taken together, this data demonstrates that Rab11b enhances brain metastasis formation following lodging in the brain parenchyma through specific interactions with the brain microenvironment, and loss of Rab11b is sufficient to decrease breast cancer brain metastasis.

**Figure 3.**
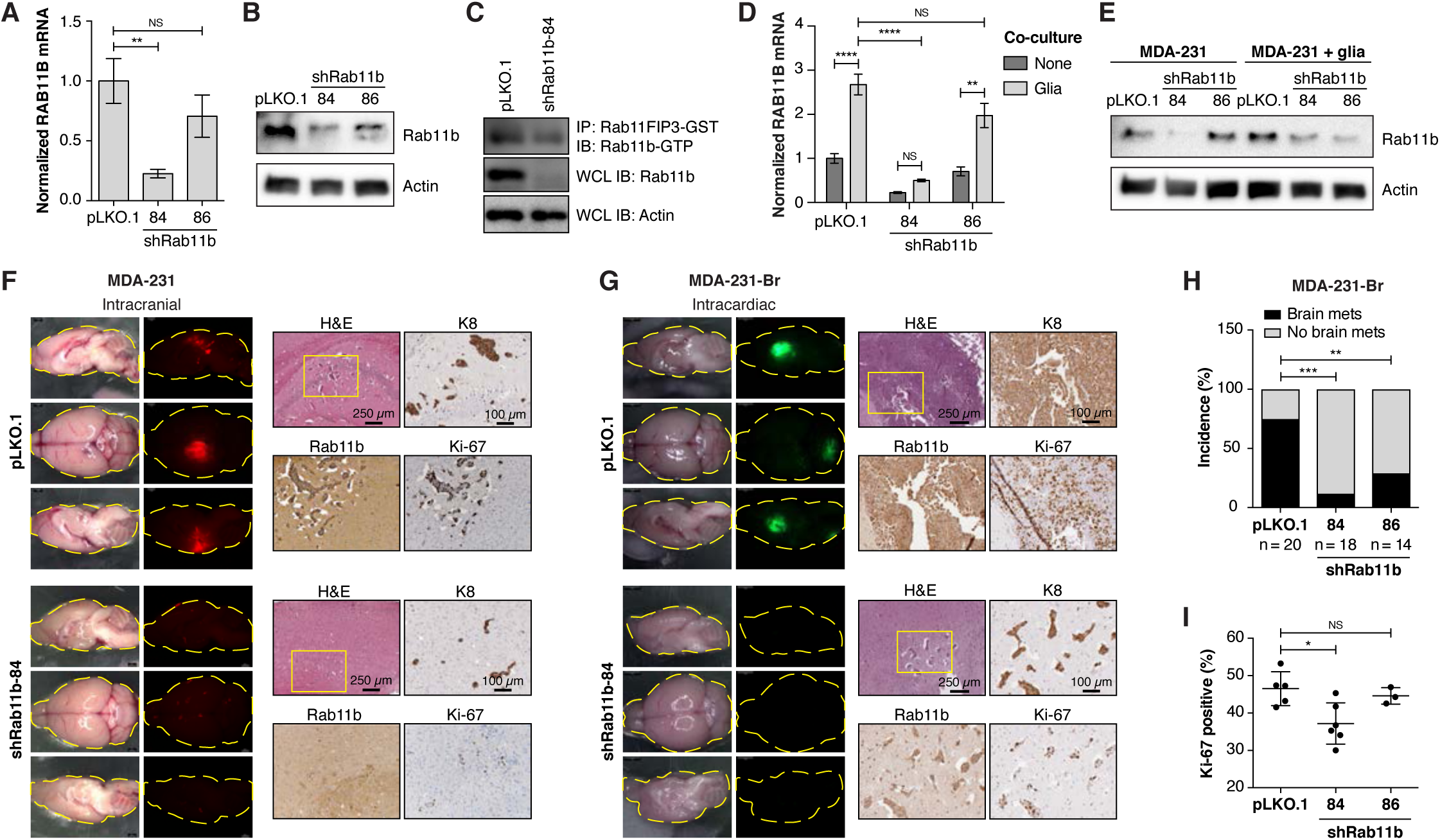
Rab11b is required for breast cancer brain metastasis. **(A)** Normalized mean RAB11B expression in MDA-231 cells, relative to pLKO.1 empty vector. ANOVA with Dunnett’s multiple comparison test. **(B)** Rab11b immunoblots for MDA-231 cells expressing indicated constructs. **(C)** Rab11b-GTP immunoblot for MDA-231 cells. **(D)** Normalized RAB11B expression in MDA-231 cells cultured alone or with primary murine glia for three days, relative to pLKO.1 alone. Two-way ANOVA with Tukey’s multiple comparison test. **(E)** Rab11b immunoblots for MDA-231 cells cultured alone or with primary murine glia for five days followed by removal of glial cells. **(F)** Representative H&E and IHC images for mice intracranially injected with MDA-231-tdTomato control or shRab11b cells. **(G)** Representative H&E and IHC images for mice intracardially injected with MDA-231-Br-EGFP control or shRab11b cells. **(H)** Incidence of brain metastasis determined by visible GFP signal at 28 dpi. Analysis of contingency, Fisher’s exact test. **(I)** Quantitation of Ki-67 staining from MDA-231-Br brain metastases. ANOVA with Dunnett’s multiple comparison test. For all panels, * p < 0.05, ** p < 0.01, *** p < 0.001, **** p < 0.0001.

### Rab11b modulates the cell surface proteome by controlling cellular recycling

Given the role of Rab11b in controlling cell surface protein recycling, we first examined Rab11b-mediated recycling of the transferrin receptor, a canonical Rab11 cargo protein. Transferrin receptor recycling is unique because transferrin and transferrin receptor remain bound during internalization and recycling to the surface, dissociating only upon being returned to the cell surface (Bleil and Bretscher, 1982; Dautry-Varsat et al., 1983), allowing monitoring of internalization and recycling. Control or shRab11b cells were labeled with fluorescent transferrin, and the rates of internalization and recycling were determined by monitoring retention of transferrin-Alexa647. Compared with control cells, severe loss of Rab11b (shRab11b-84, Figure 3B) led to decreased recycling (Figure 4A, Figure S5A), while internalization was not affected (Figure S5B), consistent with the role of Rab11b in regulating transport of proteins from the endosomal recycling center (ERC) to the cell surface (Grant and Donaldson, 2009). Interestingly, although co-culture with primary glia induces expression of Rab11b, there is not a detectable increase of transferrin receptor recycling when cells are cultured with either primary glia or CAF cells (Figure 4B, Figure S5C), suggesting that brain-mediated up-regulation of Rab11b does not globally alter the recycling of all Rab11b cargo proteins.

**Figure 4.**
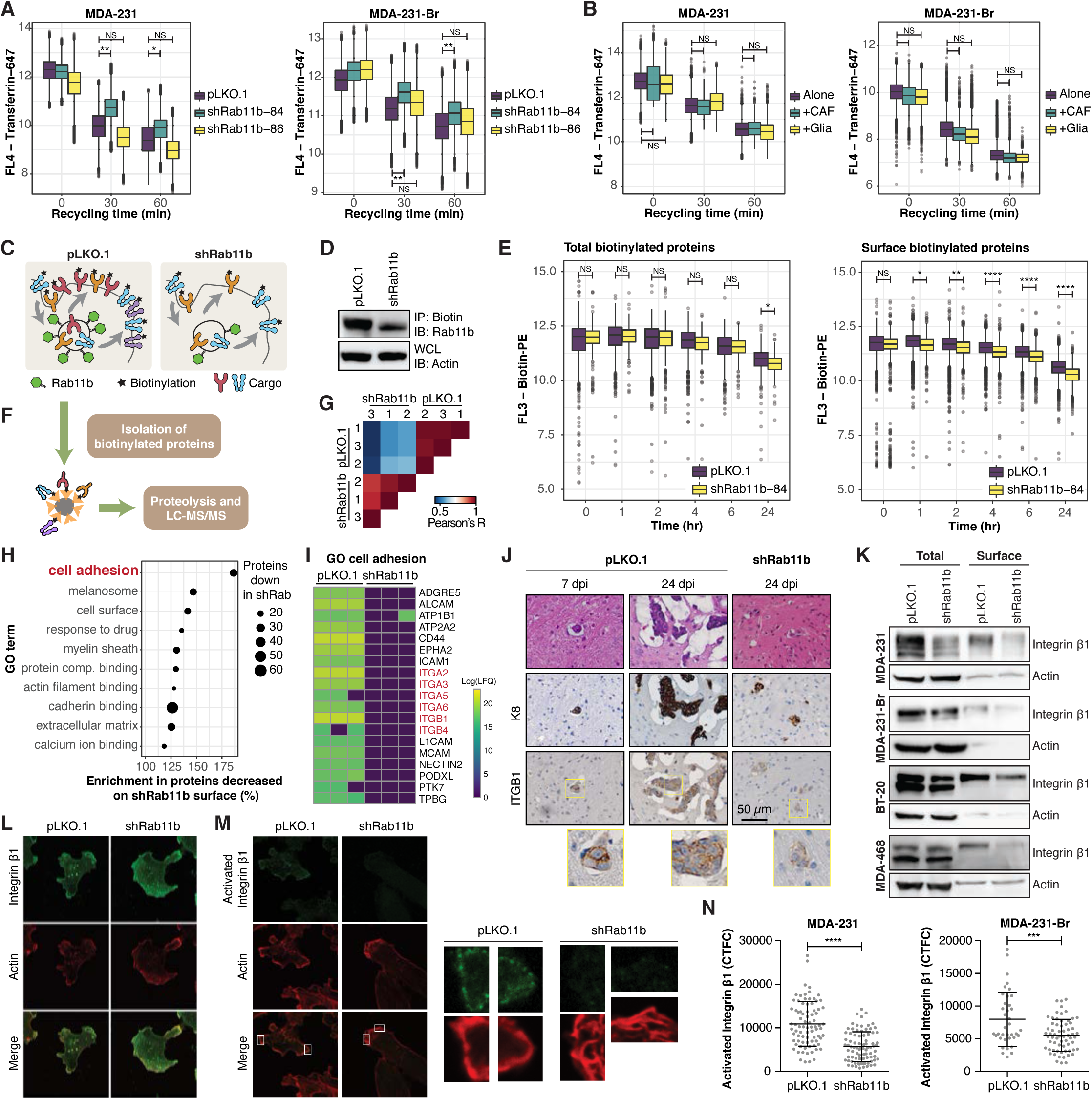
Rab11b-mediated recycling controls the cell surface proteome. **(A)** Transferrin receptor recycling in MDA-231 and MDA-231-Br cells. Two-way ANOVA with Sidak’s multiple comparison test. **(B)** Transferrin receptor recycling in cells co-cultured for two days. Cancer cells were selected on expression of CD-44. Two-way ANOVA with Sidak’s multiple comparison test. **(C)** Schematic of surface biotinylation. **(D)** Association of Rab11b with biotinylated surface proteins determined by immunoprecipitation and immunoblotting. **(E)** Retention of total or surface biotin following surface biotinylation with biotin-PE. Two-way ANOVA with Sidak’s multiple comparison test. **(F)** Schematic of biotinylated surface protein isolation and proteomics. **(G)** Correlation matrix of all measured samples based on Pearson’s correlation values. **(H)** Cleveland plot of top GO terms enriched in proteins that were decreased on the surface of shRab11b cells. **(I)** Heatmap of proteins annotated with GO term cell adhesion, containing at least one predicted transmembrane domain. **(J)** Representative images of H&E and IHC for mice intracranially injected with MDA-231-tdTomato cells. IHC for cytokeratin 8 (K8), Rab11b and integrin β1 (ITGB1). **(K)** Immunoblotting of total and surface lysates. **(L-N)** MDA-231 cells stained for integrin β1 and actin (phalloidin, to delineate cell boundaries) (L), or active integrin β1 and actin (M). (N) Corrected total cellular active integrin β1 fluorescence (CTCF) was determined for individual cells. Two-sided t-test. For all panels, * p < 0.05, ** p < 0.01, *** p < 0.001, **** p < 0.0001.

To examine Rab11b-regulated retention and recycling of surface proteins, we biotinylated cell-surface proteins with membrane-impermeable sulfo-NHS-SS-biotin (Figure 4C). Rab11b associates with internalized surface proteins, with shRab11b cells showing decreased association (Figure 4D). There was a small decrease in the retention of total biotinylated proteins in shRab11b cells (Figure 4E, left), but a significant decrease in surface expression of biotinylated proteins (Figure 4E, right), suggesting an inability of shRab11b cells to return a fraction of internalized surface proteins back to the surface. To identify the subset of surface proteins that are dependent on Rab11b for recycling, we performed mass spectrometry analysis of the cell surface proteomes of control and shRab11b cells (Figure 4F, red and purple proteins). We observed a strong correlation between biological replicates (Figure 4G). Although equal amounts of surface protein were loaded (Figure S6A), the number of proteins identified in shRab11b cells was lower (Figure S6B), approximately 200 proteins for shRab11b cells versus over 700 proteins for control cells, suggesting that there is decreased abundance and diversity in the cell surface proteome of cells without Rab11b. We employed bioinformatic analysis to focus on proteins that are known to be surface localized (through annotation with the GO slim term “Plasma Membrane”), leaving 226 proteins in pLKO.1, and 64 proteins in shRab11b cells (Figure S6C). To identify protein functional groups, the incidence of repeated GO terms was determined for proteins decreased or lost from the surface of shRab11b cells, which revealed that proteins involved in cell adhesion are dramatically decreased in shRab11b cells (Figure 4H). Further analysis of proteins involved in cell adhesion, combined with the identification of proteins with transmembrane domains using the prediction algorithm TMHMM (Krogh et al., 2001; Sonnhammer et al., 1998), revealed that integrins in particular are dramatically decreased on the surface of shRab11b cells (Figure 4I).

Integrins are heterodimeric proteins that mediate cell attachment to the extracellular matrix (ECM) (Moreno-Layseca et al., 2019). Of the integrins identified in our cell surface proteome screen, integrin β_1_ (ITGB1) is the most versatile (Humphries et al., 2006), recognizing a range of ECM proteins, and forming heterodimers with the majority of α integrins, including all of the α integrins we found to be decreased on the surface of shRab11b cells (Figure 4I). We found that integrin β_1_ is strongly expressed in brain metastases, with reduced expression in shRab11b brain metastases (Figure 4J). Examination of breast cancer cells in culture reveals that knocking down Rab11b slightly decreases overall expression of integrin β_1_, but leads to dramatic loss of cell surface integrin β_1_ (Figure 4K). Next, we examined total and active integrin β_1_ and found that while total integrin β_1_ expression was not altered in shRab11b cells (Figure 4L), there was almost a complete loss of activated integrin β_1_ (Figure 4 M, N). Taken together, this data suggests that Rab11b-mediated recycling controls the cell surface proteome, maintaining integrin β_1_ surface localization, where it can be activated by the extracellular matrix to facilitate tumor cell survival.

### Rab11b recycling of integrin β1 is necessary for tumor cell survival in the brain microenvironment

Successful ECM engagement is required for the survival of cancer cells, particularly metastatic cancer cells as they adapt to a new metastatic microenvironment (Hamidi and Ivaska, 2018). The loss of surface integrin β_1_ suggests that shRab11b cells would exhibit decreased adhesion to integrin β_1_ ligands. Indeed, we found that loss of Rab11b led to delayed ECM engagement (Figure 5A), although they were ultimately able to adhere (Figure S7A). To determine the extent of reliance on integrin β_1_, we plated cells on poly L-lysine (pLL), which does not induce integrin activation, or Type I collagen, an integrin β_1_ ligand. Control and shRab11b cells remain rounded on pLL, although control cells are better able attach and spread (Figure 5B). On collagen, control cells successfully spread and assume their characteristic mesenchymal-like morphology, while shRab11b cells showed an evident defect in spreading with reduced cell area and length (Figure 5B). Suggesting that loss of integrin β_1_ from the surface of shRab11b severely reduces cell spreading, and further that control cells possess a weak, non-integrin β_1_-mediated mechanism for spreading, which is likewise lost in shRab11b cells.

**Figure 5.**
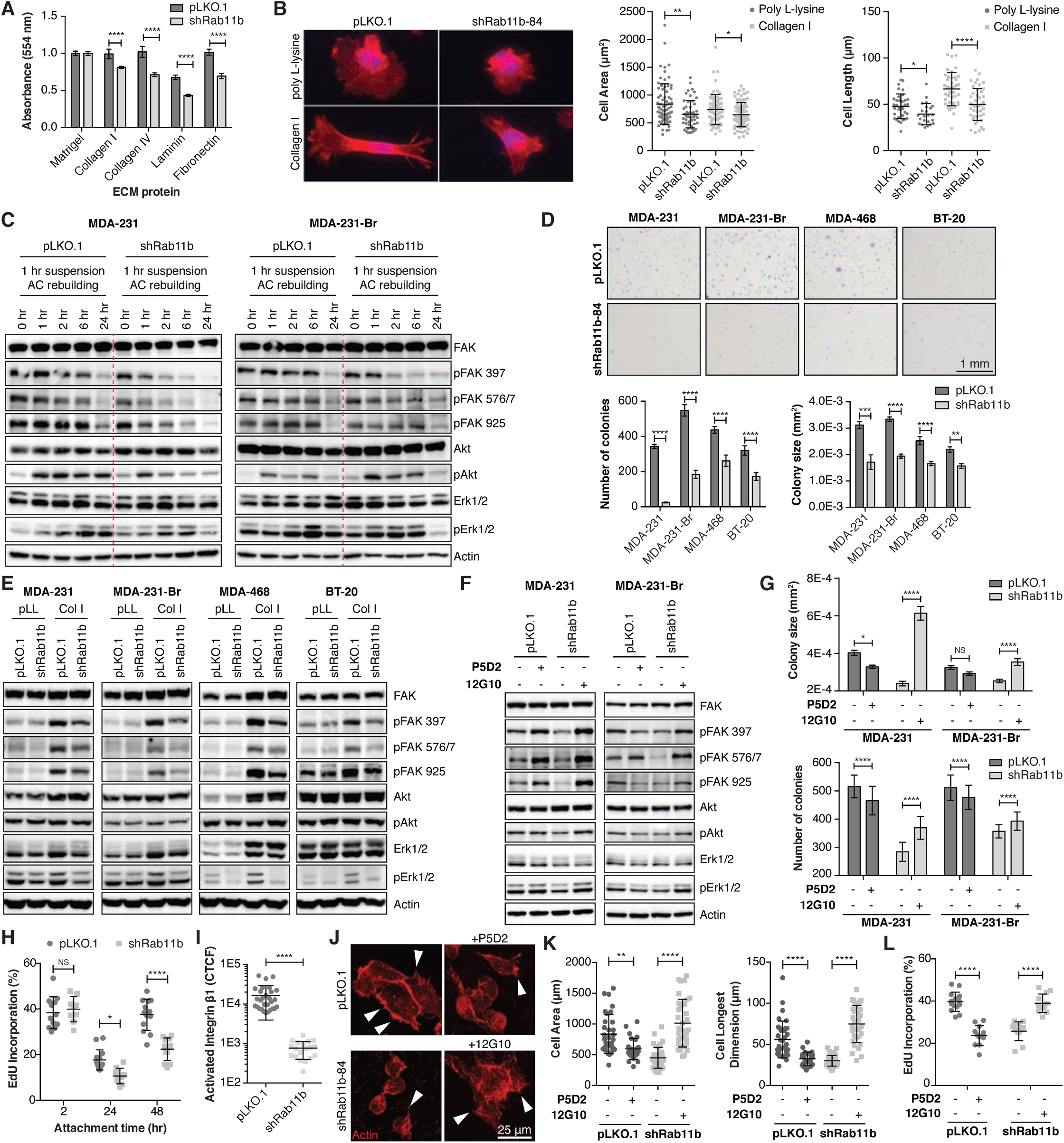
Rab11b recycling of integrin β1 is necessary for survival in the brain microenvironment. **(A)** Cell attachment at 6 hrs. Each cell line normalized to Matrigel control. Two-way ANOVA with Sidak’s multiple comparison test. **(B)** Representative images of actin (phalloidin, red) and nuclei (DAPI, blue) for cells plated on poly L-Lysine or Collagen I for 24 hrs. *Right*, quantification of cell area and longest dimension. ANOVA with Tukey’s multiple comparison test. **(C)** Cells were suspended for 1 hr at 37C, then plated to allow adhesion complex (AC) formation. Immunoblots showing signaling during adhesion. **(D)** *Top*, representative images of cells grown in soft agar for three weeks. *Bottom*, quantification of colony number and size for ten fields per condition. Two-way ANOVA with Sidak’s multiple comparison test. **(E)** Immunoblots showing signaling after 6 hr adhesion to poly L-lysine (pLL) or Collagen I (Col I). **(F)** Immunoblots showing signaling after 1 hr incubation with P5D2 (integrin β1 blocking antibody) or 12G10 (integrin β1 activating antibody), followed by 6 hr adhesion to Col I. **(G)** Quantification of colony number and size for cells treated with P5D2 or 12G10 and grown in soft agar for two weeks. Ten fields per condition. Two-way ANOVA with Sidak’s multiple comparison test. **(H)** Quantification of EdU incorporation for cells adhering to decellularized brain matrix. Ten fields per condition. Two-way ANOVA with Sidak’s multiple comparison test. **(I)** Corrected total cellular active integrin β1 fluorescence (CTCF). Two-sided t-test. **(J-L)** (J) Representative images of actin (phalloidin, red) protrusions (arrowheads) for cells treated with P5D2 or 12G10 and allowed to adhere to decellularized brain matrix for 48 hrs. (K) Actin staining was used to quantify cell area and longest dimension. ANOVA with Tukey’s multiple comparison test. (L) Quantification of EdU incorporation. Ten fields per condition. ANOVA with Tukey’s multiple comparison test. For all panels, * p < 0.05, ** p < 0.01, *** p < 0.001, **** p < 0.0001.

To examine signaling, we suspended cells for 1 hr to force down-regulation of adhesion-mediated signaling, and found that reduced Rab11b induced a severe defect in the cell’s ability to initiate and sustain attachment-mediated downstream signaling (Figure 5C). In contrast to sustained activation of FAK up to 6 hours, there was diminished phosphorylation of FAK, Erk and Akt in shRab11b cells as early as two hours post-plating (Figure 5C). Although this reduction in adhesion-mediated signaling does not change the proliferation of shRab11b cells in culture (Figure 7SB), loss of Rab11b dramatically decreases both colony size and number when cells are grown in soft agar (Figure 5D). This data suggests that the adhesion defects mediated by loss of Rab11b are masked when cells are grown in idealized cell culture conditions, likely due to the eventual ability of cells to adhere (Figure S7A), but become apparent when cells are exposed to microenvironmental stress. When cells are plated on pLL, although both lines have low activation, shRab11b cells exhibit decreased activation of FAK, Akt and Erk (Figure 5E), consistent with their decreased ability to spread (Figure 5B). Control cells dramatically increase activation of FAK, Erk and Akt when plated on Col I, suggesting functional cell surface integrin β_1_ for ligation to Type I collagen. shRab11b are not able to induce signaling to the same extent, suggesting that loss of surface integrin β_1_ induces a defect in ligation to Col I and decreased adhesion-mediated signaling (Figure 5E). To further investigate the role of Rab11b in mediating integrin β_1_ signaling, we treated control cells with an inhibitory integrin β_1_ antibody (P5D2), or shRab11b cells with an activating antibody (12G10) (Figure 5F). Inhibition of integrin β_1_ in control cells induces auto-phosphorylation of FAK, but decreases activation of Akt and Erk (Figure 5F), as expected. Activation of integrin β_1_ in shRab11b cells increases FAK, Akt and Erk signaling (Figure 5F). Furthermore, modulation of Integrin β_1_ activation status also leads to suppression (P5D2), or induction (12G10) of growth in soft agar in control and shRab11b cells, respectively (Figure 5G, Figure S7C). Together, this data suggests that Rab11b-mediated recycling of integrin β_1_ enhances the ability of breast cancer cells to successfully active attachment-mediated signaling and ultimately survive in sub-optimal ECM conditions.

As cells metastasize, they are exposed to new ECM conditions, and their ability to successfully engage this foreign ECM is critical for metastatic success. To determine whether Rab11b directly impacts the ability of breast cancer cells to engage the brain ECM, we cultured tumor cells on decellularized murine brain ECM (De Waele et al., 2015). At 2 hrs the cells have begun to adhere, and the equal proliferative rates of control and shRab11b cells in culture are apparent; however, as cells require adhesion-mediated signaling in response to the brain ECM, shRab11b cells are unable to initiate (24 hr) and sustain (48 hr) cell proliferation (Figure 5H). Immunostaining reveals considerably decreased active integrin β_1_ in shRab11b cells plated on brain ECM (Figure 5I, Figure S7D). Phalloidin staining reveals that control cells are able to spread and form protrusions on the brain matrix, while shRab11b cells remain rounded up, and activation (P5D2) or inhibition (12G10) of integrin activity prevents or induces spreading, respectively (Figure 5J, K). Inhibiting integrin β_1_ signaling prevents proliferation in control cells, while activation induces proliferation of shRab11b cell on brain matrix (Figure 5L). Taken together, this data suggests that Rab11b is required for surface localization of integrin β_1_, which in turn initiates focal adhesion signaling, allowing attachment to the brain ECM and ultimately survival in the brain metastatic microenvironment.

### Inhibition of Rab11b function decreases breast cancer brain metastasis

Given the ability of Rab11b to control the cell surface proteome, including recycling of integrin β_1_, we sought to pharmacologically inhibit Rab11b. Historically, targeting small GTPases has not been easily accomplished or translated (Brock et al., 2016; Cox et al., 2015; Pei et al., 2018); however, a unique feature of Rab proteins is that they require a lipid modification that allows membrane localization, which is required for function (Joberty et al., 1993). Specifically, Rab proteins are geranylgeranylated, and the geranylgeranylpyrophosphate required is generated by the mevalonate pathway (Figure 6A) (Resh, 2013). The mevalonate pathway is a inhibited by HMG-CoA reductase inhibitors such as statins (Goldstein and Brown, 1990). Thus, we postulate that statin treatment could suppress Rab11b activation, an essential step for breast cancer adaptation to the brain metastatic microenvironment. Because brain metastases, particularly early brain metastases, are protected by the blood-brain barrier (Lockman et al., 2010; Palmieri et al., 2006b), we chose two lipophilic statins, pitavastatin (Pit) and simvastatin (Sim), which are reported to be blood-brain barrier permeable (Ifergan et al., 2006; Morofuji et al., 2010). We found that both statins successfully inhibited tumor cell growth in soft agar (Figure 6B), phenocopying the effects of Rab11b knockdown (Figure 5D). Geranylgeranylation is required for Rab11b membrane association, and pitavastatin or simvastatin treatment moved Rab11b from the insoluble, membrane bound fraction to the soluble, non-membrane bound fraction in a dose dependent manner (Figure 6C, Figure S8A). Importantly, statin treatment also decreased the amount of active Rab11b (Figure 6D), and decreased the rate of transferrin receptor recycling (Figure 6E), suggesting that both statins effectively inhibit geranylgeranylation of Rab11b, thereby preventing localization and function.

**Figure 6.**
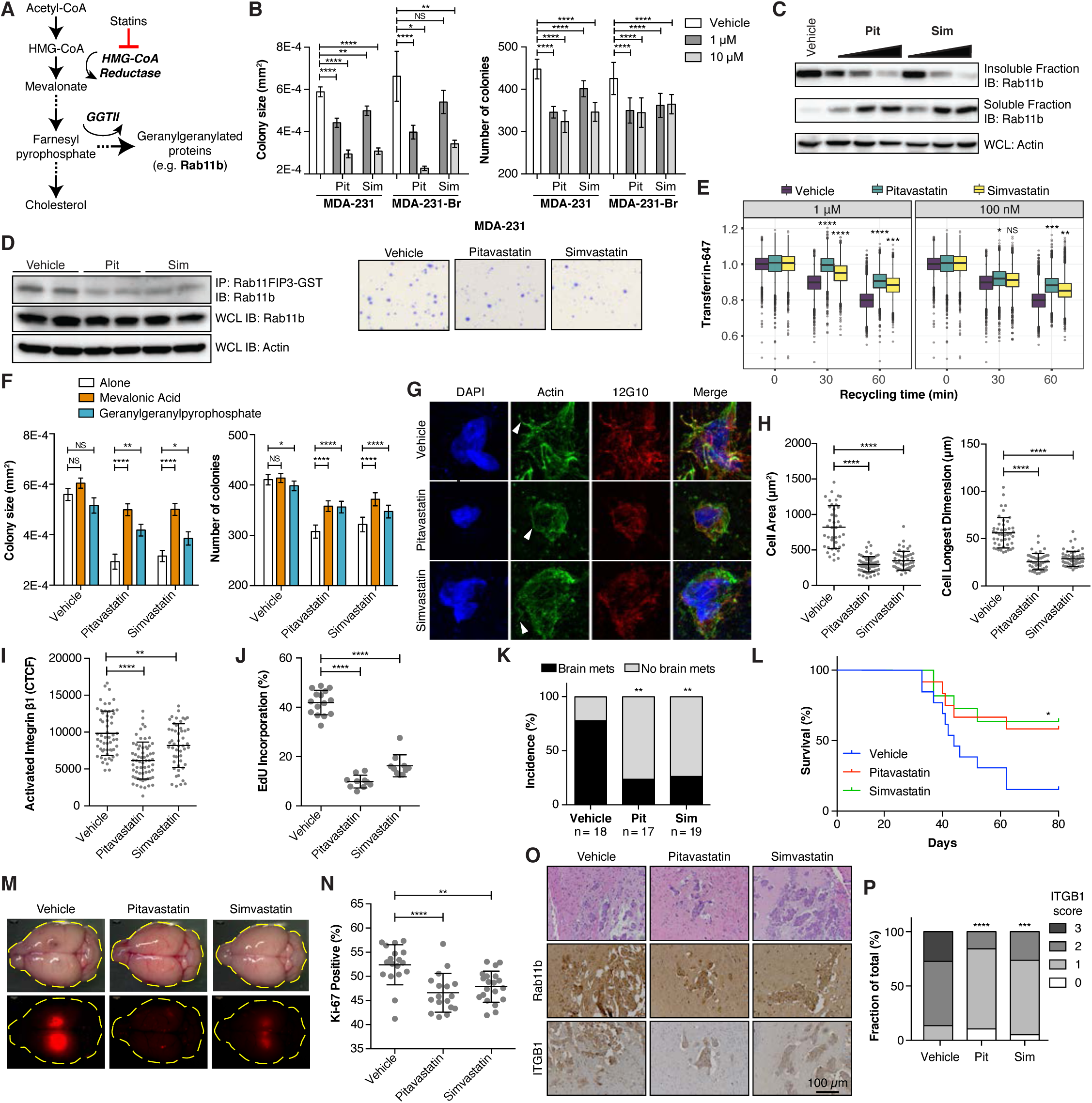
Inhibition of Rab11b function decreases breast cancer brain metastasis. **(A)** Schematic of the mevalonate pathway leading to Rab11b geranylgeranylation. **(B)** Quantification of number and size of colonies for cells grown in soft agar for three weeks in the presence of vehicle or 1 μM pitavastatin or simvastatin. *Bottom*, representative images. Two-way ANOVA with Sidak’s multiple comparison test. **(C)** MDA-231 cells grown with vehicle or 10 μM-100 nM pitavastatin or simvastatin for 24 hrs. Immunoblotting of soluble and insoluble fractions separated with Triton X-114. **(D)** Rab11b activation assay for MDA-231 cells grown with of vehicle or 1 μM pitavastatin or simvastatin for 24 hrs, followed by immunoblotting. **(E)** Transferrin receptor recycling in MDA-231 cells grown with vehicle or pitavastatin or simvastatin for 24 hrs. Two-way ANOVA with Sidak’s multiple comparison test. **(F)** Quantification of colony size and number for MDA-231 cells grown in soft agar for two weeks with 1 μM pitavastatin or simvastatin, with the addition of 100 μM mevalonic acid or 10 μM geranylgeranylpyrophosphate. Ten fields per condition. Two-way ANOVA with Sidak’s multiple comparison test. **(G-J)** MDA-231 cells allowed to adhere to decellularized murine brain matrix for 48 hrs in the presence of 1 μM pitavastatin or simvastatin. (G) Representative images of actin (phalloidin, green) protrusions (arrowheads), active integrin β1 (12G10, red), and nuclei (DAPI, blue). (H) Quantification of cell area and longest dimension. ANOVA with Tukey’s multiple comparison test. (I) Corrected total cellularactive integrin β1 fluorescence (CTCF). ANOVA with Tukey’s multiple comparison test. (J) Quantification of EdU in ten fields per condition. ANOVA with Tukey’s multiple comparison test. **(K-L)** MDA-231-Br-GFP cells were intracardially injected, and given daily intraperitoneal injections of vehicle or 1 mg/kg pitavastatin or 5 mg/kg simvastatin. (K) Incidence of brain metastasis determined by visible GFP signal at 28 dpi. Analysis of contingency. (L) Survival determined by daily monitoring for the apearance of neurological symptoms or euthanasia criteria. Log rank test. **(M-P)** MDA-231-tdTomato cells were intracranially injected, and given daily intraperitoneal injections of vehicle or 1 mg/kg pitavastatin or 5 mg/kg simvastatin. (M) Representative images of brain metastasis at 34 dpi. (N) Quantification of Ki-67 staining for proliferation. ANOVA with Tukey’s multiple comparison test. (O) Representative images of Rab11b and ITGB1 immunostaining. (P) Scoring of ITGB1 immunostaining. Analysis of contingency. For all panels, * p < 0.05, ** p < 0.01, *** p < 0.001, **** p < 0.0001.

Statins inhibit HMG-CoA reductase, the enzyme responsible for generating mevalonic acid (Goldstein and Brown, 1990), and have broad effects beyond inhibition of Rab proteins generally, or Rab11b specifically. Pitavastatin and simvastatin have both been shown to inhibit breast cancer proliferation in vitro and in vivo by inhibiting RhoA, NF-κB, or PI3K signaling (Campbell et al., 2006; Denoyelle et al., 2001; Ghosh-Choudhury et al., 2010; Gronich and Rennert, 2013; Park et al., 2013; Wang and Kitajima, 2007). To determine if statin treatment decreases tumorigenicity in part through inhibition of Rab proteins, we performed a metabolite rescue experiment. Cells were treated with statins in addition to mevalonic acid (MVA), which restores all signaling downstream of HMG-CoA, including geranylgeranylation of Rab proteins. Restoration of the entirety of the mevalonate pathway with MVA rescues growth in soft agar as expected (Figure 6F, Figure S8B). Significantly, addition of geranylgeranylpyrophosphate, a metabolite essential for geranylgeranylation of Rab proteins, significantly rescued growth of tumor cells in soft agar, increasing both colony size and number (Figure 6F, Figure S8B), suggesting that statins mediate their effect on tumorigenicity in part through inhibition of Rab localization and function.

We next sought to determine if statins would prevent Rab11b-mediated recycling of integrin β_1_. Cells were plated on decellularized brain matrix and treated with statins for 48 hrs. Statin treatment prevents cell spreading (Figure 6G, H), decreases activation of integrin β_1_ (Figure 6I), and inhibits proliferation of breast cancer cells on the brain ECM (Figure 6J), suggesting that statins successfully inhibit Rab11b-mediated recycling of integrin β_1_. To confirm this effect in vivo, we used both intracardiac and intracranial models of breast cancer brain metastasis. Beginning two days post injection, animals given daily treatment with human equivalent doses of pitavastatin or simvastatin showed a dramatic decrease in brain metastasis incidence (Figure 6K, M), with a concurrent increase in survival (Figure 6L). Brain metastases exhibited decreased proliferation (Figure 6N, Figure S8C), Given that statins inhibit membrane localization of Rabs, statin treatment did not alter Rab11b expression as expected (Figure 6O). However, statin treatment significantly decreases the expression of integrin β_1_ in brain metastases (Figure 6O, P), suggesting that statin treatment inhibits Rab11b-mediated recycling of integrin β_1_ to suppress breast cancer brain metastasis.

## DISCUSSION

Metastatic disease is an undisputedly urgent clinical problem, and patients with brain metastases protected by the blood-brain barrier are in particular need of effective therapeutic options. We provide here two significant insights into breast cancer brain metastasis. First, whereas metastatic adaptation is often studied at the transcriptome level, our data shows that endosomal recycling can dramatically alter the ability of DTCs to interact with their microenvironment through control of protein localization to the cell surface. Several recent studies combining proteomic and genomic analysis have demonstrated a low degree of concordance between mRNA and protein expression (Johansson et al., 2019; Mertins et al., 2016; Sinha et al., 2019; Zhang et al., 2016), highlighting the importance of protein-level regulation, such as recycling. Second, we demonstrate the pre-clinical efficacy of statin treatment for BCBM, and provide a mechanistic rationale for this efficacy, through inhibition of Rab11b geranylgeranylation.

By combining temporal transcriptional analysis of breast cancer brain metastasis formation with functional screening, we identify genes that are not only transcriptionally dysregulated, but functionally driving tumorigenesis and metastasis. We show that breast cancer cells significantly up-regulate Rab11b in the brain metastatic site compared to the primary site. Previous studies have identified diverse mechanisms controlling breast cancer survival in the brain metastatic microenvironment, often relying on transcriptional profiling of cancer or microenvironmental cells (Bos et al., 2009; Park et al., 2011). Our finding that endosomal recycling exerts control over the cell surface proteome suggests that it is an important, yet previously unconsidered, level of regulation coordinating metastatic adaptation.

Specifically, we show that loss of Rab11b decreases retention of proteins on the cell surface, with a dramatic loss of several integrins, including integrin β_1_. It is well known that localization governs protein function, particularly for adhesion proteins and growth factor receptors such as E-cadherin and EGFR (Hung and Link, 2011; Pellinen et al., 2006; Thiery, 2002; Ye et al., 2016). We find that Rab11b-mediated recycling of integrin β_1_ controls surface expression, and therefore ECM ligation, leading to decreased attachment and spreading on integrin β_1_ ligands. Integrin β_1_ is the most versatile β isoform (Humphries et al., 2006), with the ability to form heterodimers with a variety of α isoforms. Rab11b-mediated loss of surface integrin β_1_ leads to decreased activation of adhesion-mediated survival signaling, rendering cells more sensitive to ECM composition. Rab11b knockdown cells are unable to spread and proliferate on decellularized brain matrix, consistent with our finding that loss of Rab11b dramatically reduces brain metastasis formation. Activation of integrin β_1_ restores attachment and survival of Rab11b cells, suggesting that forced clustering and activation of integrin β_1_ is able to overcome decreased recycling. Although recycling-mediated localization of specific proteins has been individually studied before, our study demonstrates that, through control of a subset of proteins, Rab11b controls the cell surface proteome to mediate breast cancer metastatic adaptation to the brain microenvironment. Thus, we propose that the Rab11b-regulated subset of proteins whose expression and localization are dictated by endosomal recycling should be considered the “recycleome”, an additional layer of control over cellular behavior. Although we found that Rab11b is the specific isoform up-regulated during BCBM, it is likely that the specific combination of primary cancer type and metastatic microenvironment will dictate the regulation and content of the recycleome. Indeed, the dependence on Rab proteins and effectors has been shown to vary from 2D to 3D culture (Mrozowska and Fukuda, 2016), highlighting the importance of studying trafficking in a specific tumor microenvironmental context.

Breast cancer brain metastases often exhibit a long latent period (Steeg, 2016), and it would be possible to target this critical adaptation period to subsequently prevent metastatic outgrowth. Given the importance of Rab11b in mediating adaptation to the brain metastatic microenvironment, we sought a therapeutic strategy to target Rab11b. The requirement for geranylgeranylation of Rab11b for function provides a unique opportunity to target the mevalonate pathway for non-specific Rab11b inhibition. The Rab family of GTPases encompasses over 70 family members, with each member localizing to distinct, but overlapping membranes within the cell (Barr, 2013). However, all Rab proteins require geranylgeranylation, suggesting that inhibition of the mevalonate pathway and its downstream geranylgeranylation with statins could be extended to multiple Rab-mediated clinical scenarios, such as prevention of brain metastasis. Long-term chemoprevention for potential brain metastasis relapse requires the proposed chemopreventive agent to be inexpensive, efficacious, and most importantly, with minimal adverse effects (Achrol et al., 2019; Steeg, 2016). The well-tolerated HmG-COA reductase inhibiting statins are ideal candidates for direct drug repurposing for brain metastasis prevention. Given the prevalence of statin use, clinical studies into the effect of statins on cancer have been generally confined to clinical meta-analysis (El-Refai et al., 2017; Gronich and Rennert, 2013). Although the mechanism for statins in cardiovascular disease is well established, the role of statins in cancer treatment is not well defined. Despite variations among different clinical studies, the general consensus is that statins, particularly lipophilic statins, are beneficial to patients with breast cancer (Ahern et al., 2011, 2014; Manthravadi et al., 2016). A number of studies have demonstrated the anti-cancer activity of statins in various pre-clinical cancer models and clinical studies (Campbell et al., 2006; Denoyelle et al., 2001; Ghosh-Choudhury et al., 2010; Park et al., 2013; Wang and Kitajima, 2007). In this study, we further provided a strong pre-clinical rationale (Figure 6), based on Rab11b-mediated metastatic adaptation mechanisms, that repurposing statins could be a clinically practical strategy for brain metastasis prevention. In conclusion, our work highlights the importance of the Rab11b-mediated recycling pathway in brain metastatic adaptation and outgrowth. Our study provides a mechanistic rationale for the use of statins in chemoprevention of breast cancer brain metastasis.

## ACKNOWLEDGEMENTS

**List of grants**

This work was partially funded by an Advancing Basic Cancer Research grant from the Walther Cancer Foundation (S.Z. and J.Z.Z.), DOD W81XWH-15-1-0021 (S.Z.) and NIH grants R01CA194697-01 (S.Z.), TL1TR001107 (E.N.H.), F32CA210583-01 (E.N.H.).

We would additionally like to acknowledge and thank the Dee Family endowment (S.Z.).

We would also like to members of the Zhang and D’Souza-Schorey labs for scientific insight and support. All sample preparation and LC/MS/MS analysis was performed at the Purdue Proteomics Facility. The Q Exactive Orbitrap HF mass spectrometer and the UltiMate 3000 HPLC system used for this study were purchased with generous funding from the Purdue Office of the Executive Vice President for Research and Partnership. We are grateful for the use of the following core facilities: Notre Dame Genomics and Bioinformatics Core Facility, Notre Dame Freimann Life Sciences Center, Indiana University School of Medicine South Bend Imaging and Flow Cytometry Core.

## AUTHOR CONTRIBUTIONS

Conceptualization, ENH, SZ

Methodology, ENH, UKA, JL

Software, ENH, MDB, PMS, ATS, JL

Formal Analysis, ENH, MDB, PMS, UKA, ATS

Investigation, ENH, MDB, MEJ, JWC, IHG, VH, UKA

Writing, Original Draft, ENH

Writing Review & Editing, ENH, SZ

Supervision, CDS, JZZ, SZ

Funding Acquisition, ENH, JZZ, SZ

---

**Figure S1.**
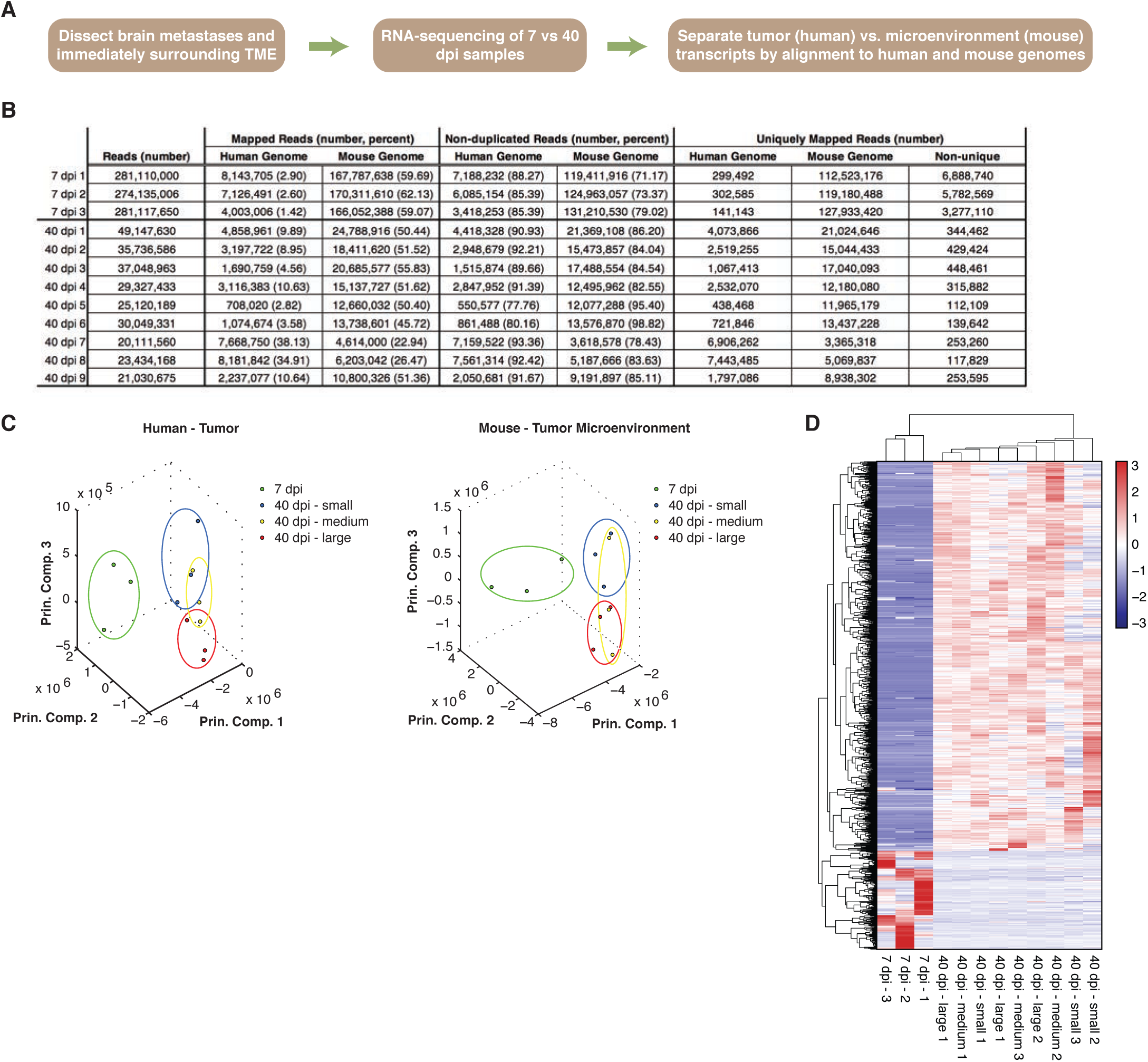
RNA-sequencing of single cell versus overt breast cancer brain metastases, Related to Figure 1. **(A)** Schematic of experimental and bioinformatic procedure. **(B)** Table of reads per sample, aligned to human or mouse genome, before and after separation based on unique alignment. **(C)** PCA analysis of human (left) and mouse (right) RNA-sequencing samples. **(D)** Heatmap of 1015 genes that were significantly differentially expressed between 7 and 40 dpi samples with a q value less than 0.05.

**Figure S2.**
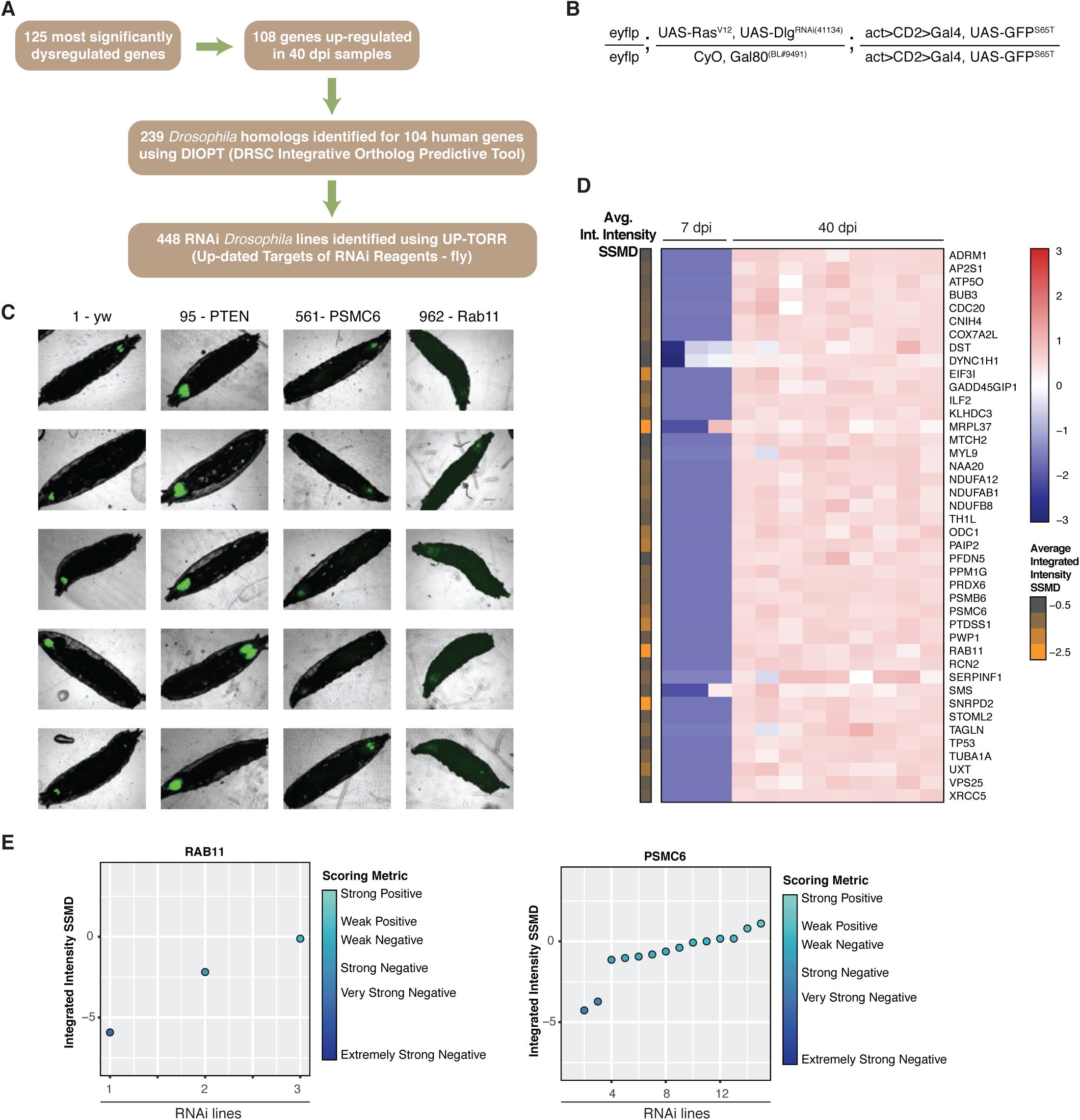
*Drosophila menalonogastor* model characterization and selection of hit genes. Related to Figure 1. **(A)** Schematic of identification of *Drosophila* orthologs and RNAi lines. **(B)** Genotype of the tumor tester line. **(C)** Representative images showing tumor signal (GFP positive, green) in larvae from negative control (yw), positive control (shPTEN), and two hit genes (PSMC6 and Rab11). **(D)** Heatmap showing expression of hit genes in original human RNA sequencing data, annotated with the average strictly standardized mean difference (SSMD) of the integrated intensity from the *Drosophila* screen. **(E)** All integrated intensity SSMD data for two representative hit genes.

**Figure S3.**
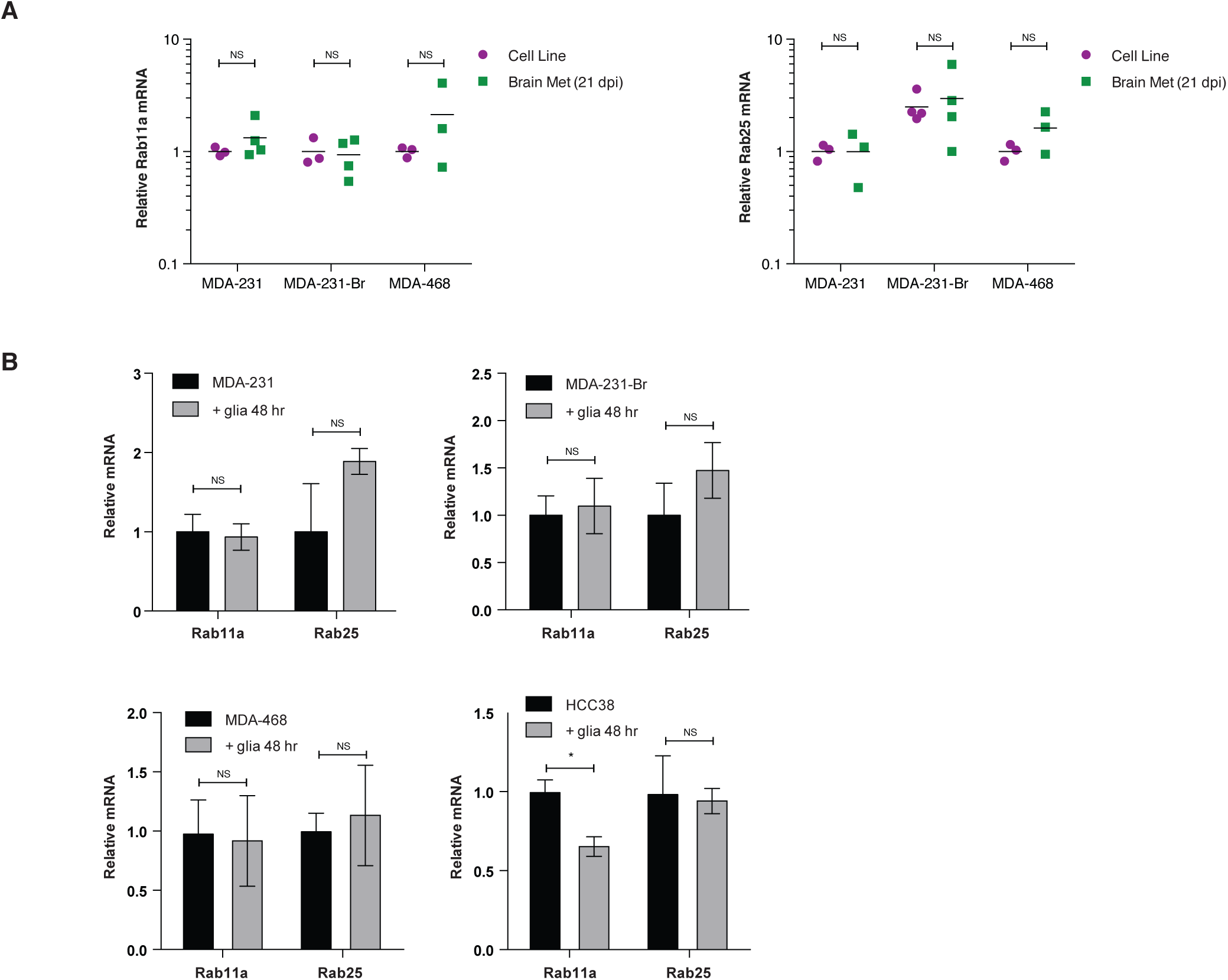
Only the Rab11 B isoform is up-regulated by the brain metastatic microenvironment. **(A)** qPCR for Rab11a and Rab25 in cells grown in culture versus brain metastases. All values are normalized to MDA-231 cells in culture. Student’s t-test. **(B)** qPCR for Rab11a and Rab25 in cell lines co-cultured with primary murine glia or CAF cells for 2 days. All values normalized to single culture For all panels, * p < 0.05.

**Figure S4.**
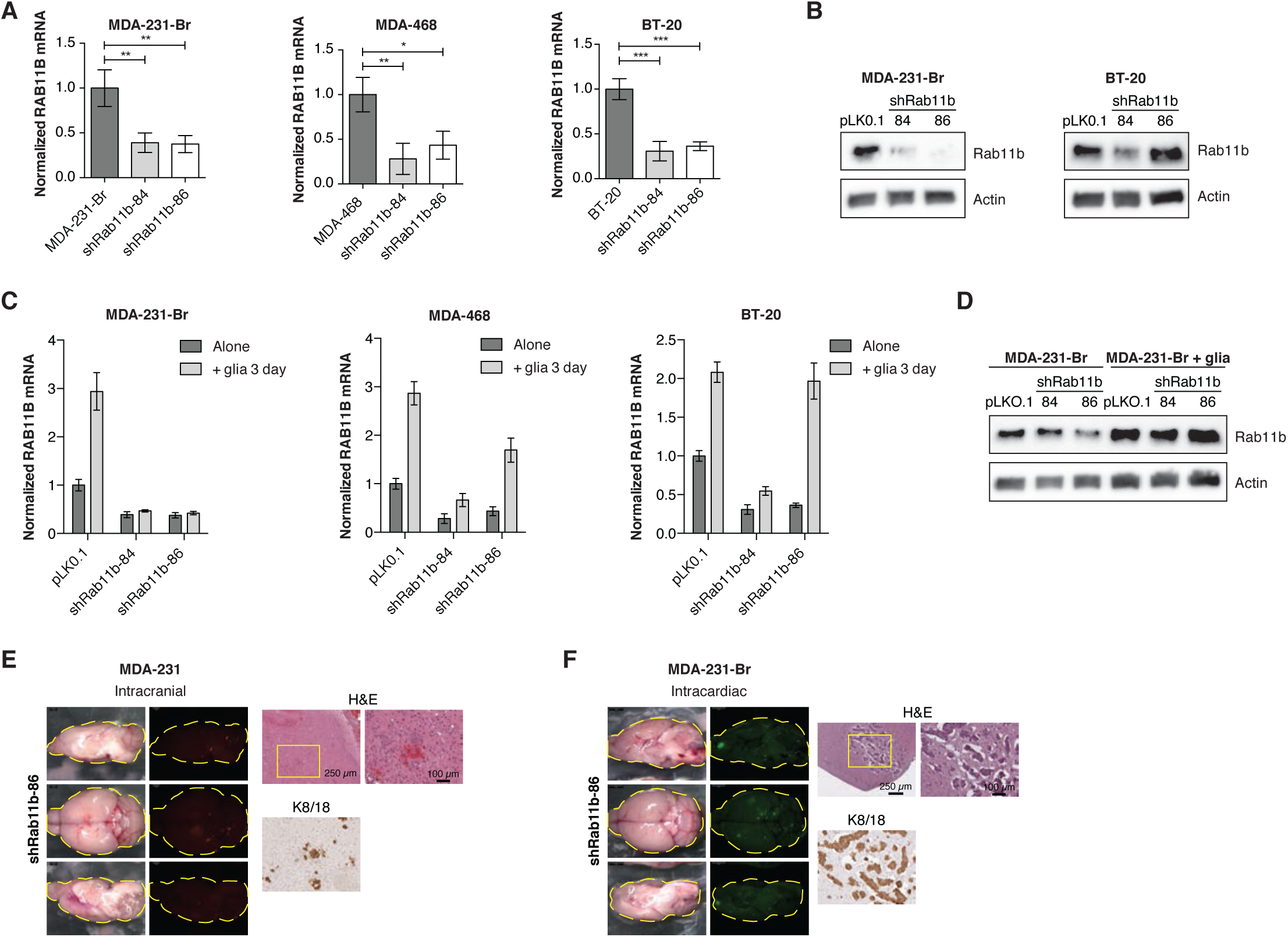
Rab11b knockdown prevents glial-mediated Rab11b up-regulation. **(A)** Normalized mean RAB11B expression in cells expressing indicated constructs, relative to pLKO.1 empty vector. ANOVA with Dunnett’s multiple comparison test. **(B)** Rab11b immunoblots for cells expressing indicated constructs. **(C)** Normalized RAB11B expression in cells expressing indicated constructs cultured alone or with primary murine glia for three days, relative to pLKO.1 alone. Two-way ANOVA with Tukey’s multiple comparison test. **(D)** Rab11b immunoblots for cells expressing indicated constructs cultured alone or with primary murine glia for five days followed by removal of glial cells using magnetic bead-based stromal cell depletion. **(E)** Representative images of mice intracranially injected with MDA-231-tdTomato shRab11b-86 cells. Mice were sacrificed 24 dpi, imaged, and H&E and immunohistological staining performed for cytokeratin 8/18 (K8/18). **(G)** Representative images of mice intracranially injected with MDA-231-Br-EGFP shRab11b-86 cells. Mice were sacrificed 28 dpi, imaged, and H&E and immunohistological staining performed for cytokeratin 8/18 (K8/18). For all panels, * p < 0.05, ** p < 0.01, *** p < 0.001, **** p < 0.0001.

**Figure S5.**
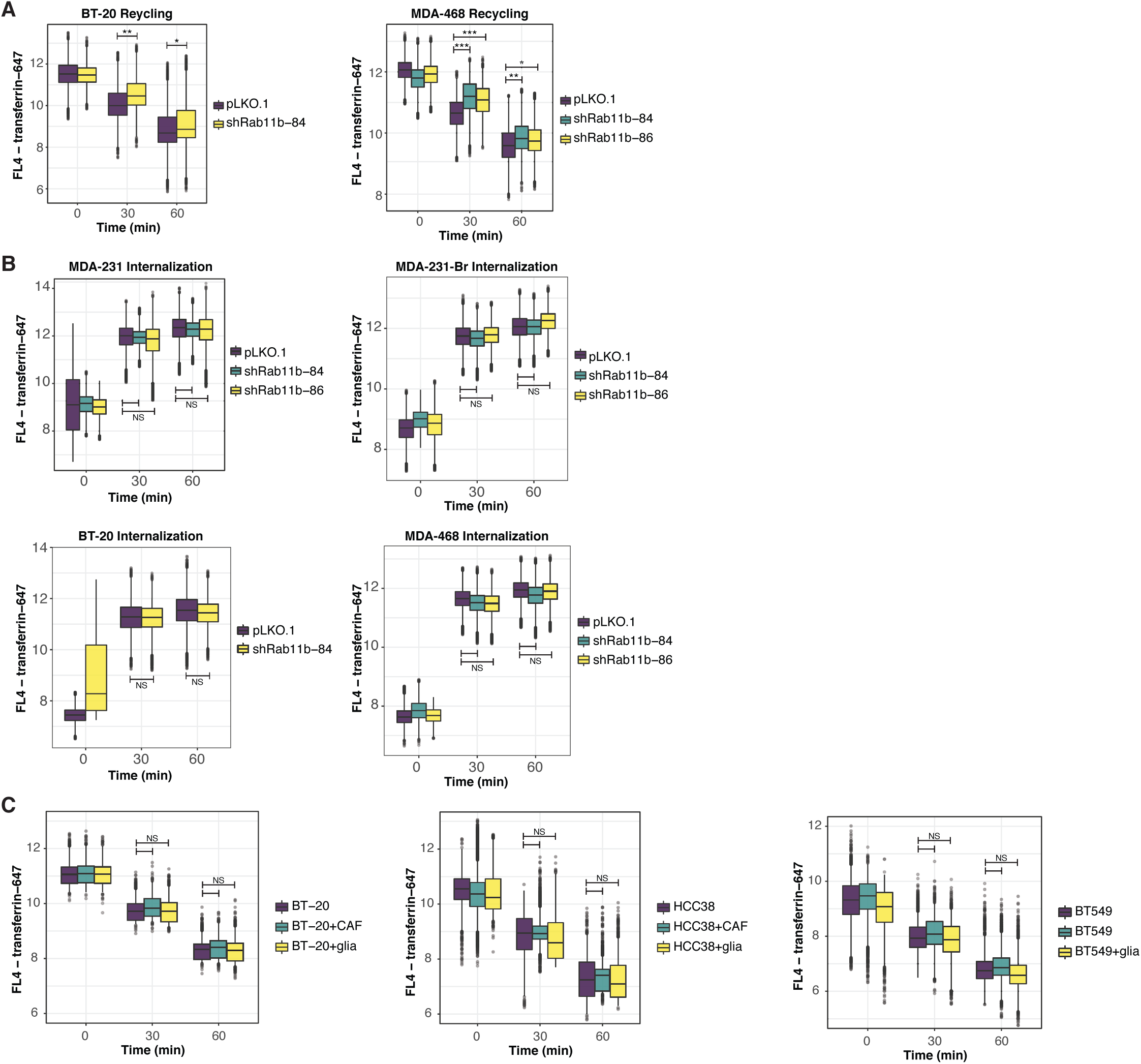
Internalization and recycling of the transferrin receptor. Related to Figure 4. **(A)** Recycling of the transferrin receptor. BT-20, ANOVA with Tukey’s multiple comparison test. MDA-468, two-way ANOVA with Sidak’s multiple comparison test. **(B)** Internalization of the transferrin receptor. MDA-231, MDA-231-Br, MDA-468, two-way ANOVA with Sidak’s multiple comparison test. BT-20, ANOVA with Tukey’s multiple comparison test. **(C)** Cells were co-cultured for two days then recycling assay performed as in A. Cancer cells were selected on expression of CD-44. Two-way ANOVA with Sidak’s multiple comparison test. For all panels, * p < 0.05, ** p < 0.01, *** p < 0.001, **** p < 0.0001.

**Figure S6.**
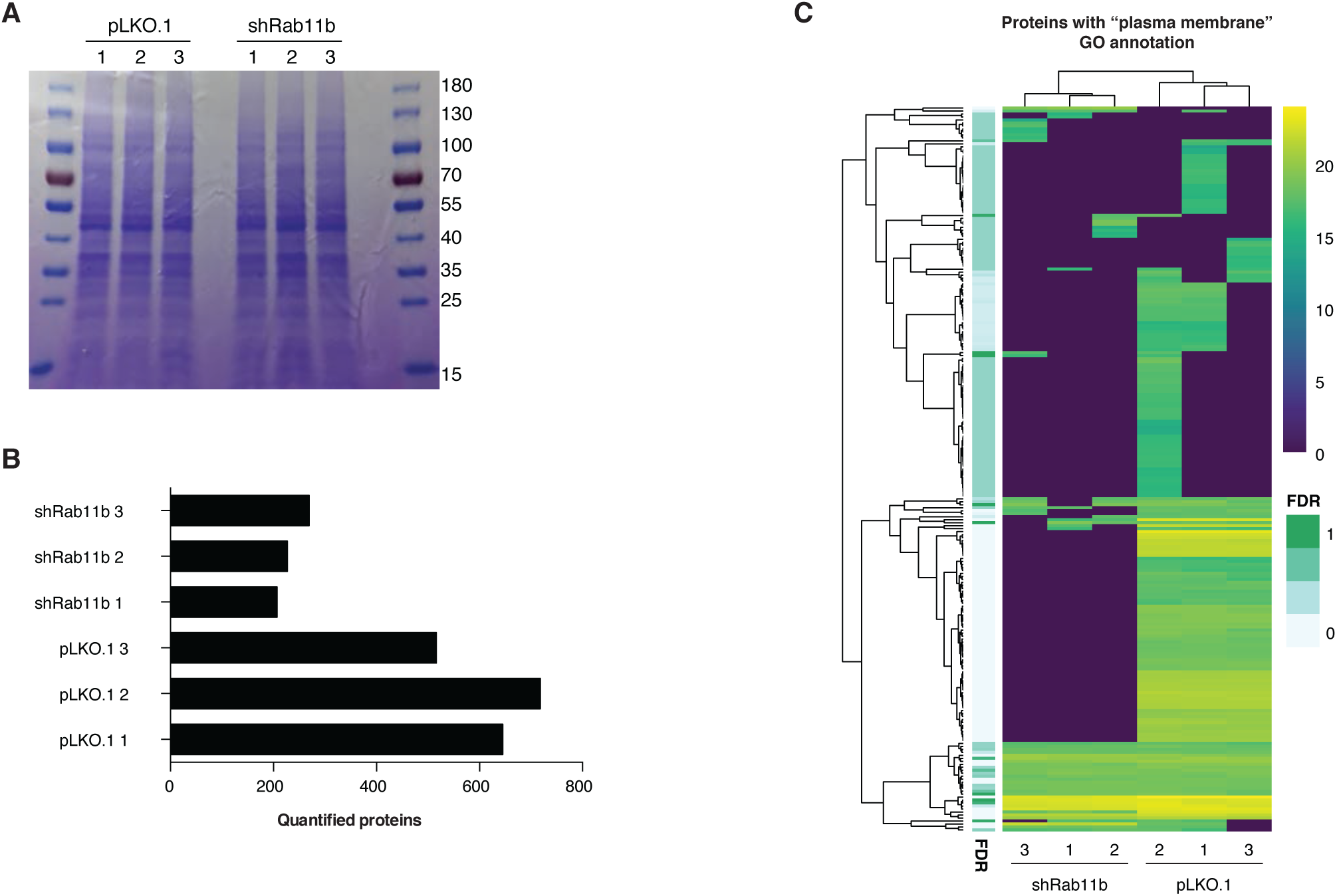
Cell surface proteome analysis. Related to Figure 4. **(A)** Surface biotinylated MDA-231 cells were immunoprecipitated, run on a protein gel and coomassie stained. **(B)** Total number of proteins identified in each sample. **(C)** Heatmap showing all identified proteins that are annotated with the GO term plasma membrane.

**Figure S7.**
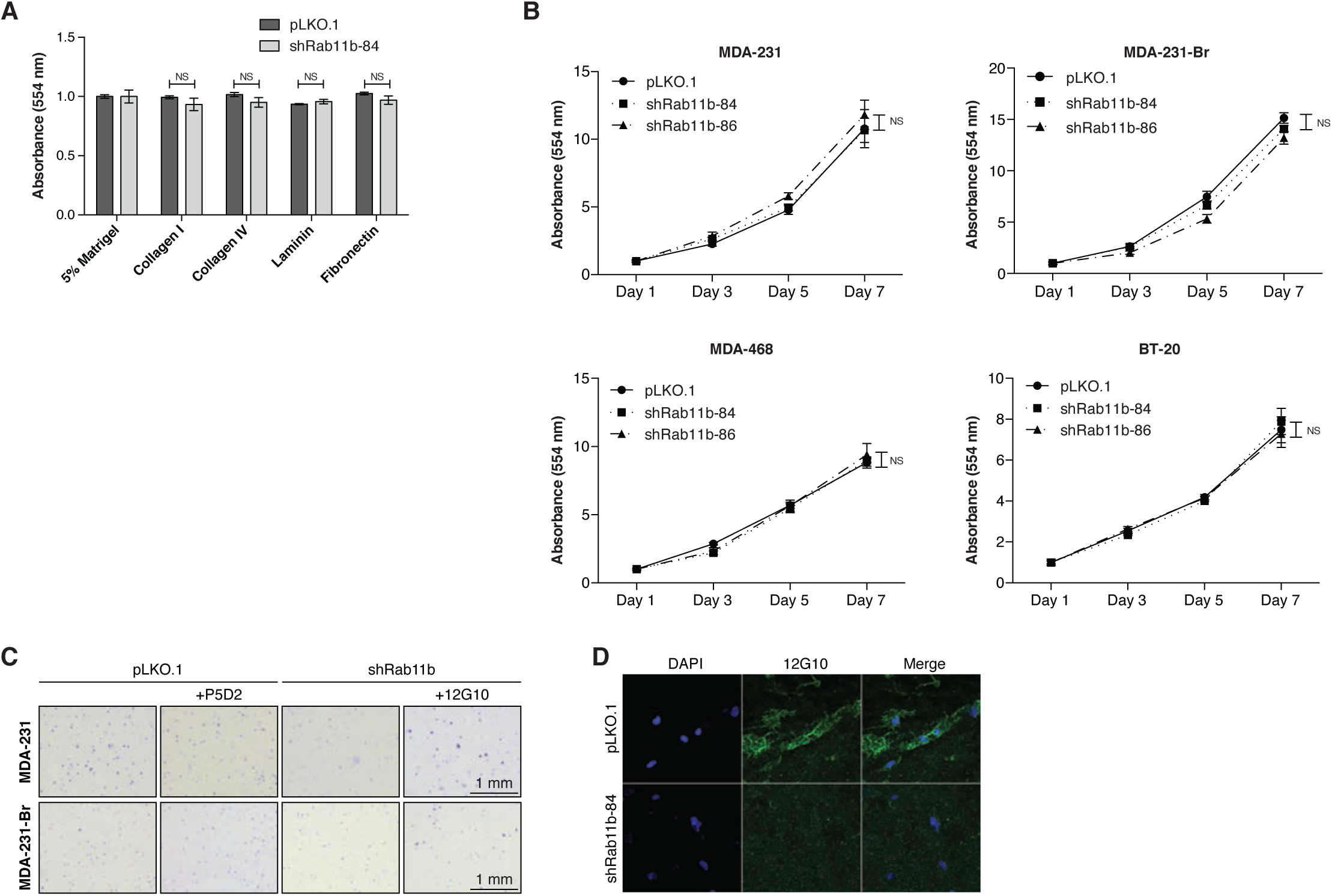
Rab11b control of adhesion and growth. Related to Figure 5. **(A)** Adhesion assay for MDA-231 cells given 48 hrs to attach. Each cell line normalized to Matrigel control. Two-way ANOVA with Sidak’s multiple comparison test. **(B)** Cell proliferation. Two-way ANOVA with Sidak’s multiple comparison test. **(C)** *Left*, Representative images of cells grown for 2 weeks in soft agar in the presence of P5D2 or 12G10. **(D)** Representative images of MDA-231 cells allowed to adhere to decellularized brain matrix for 48 hr, and stained for active integrin β1 (12G10, green), and nuclei (DAPI, blue).

**Figure S8.**
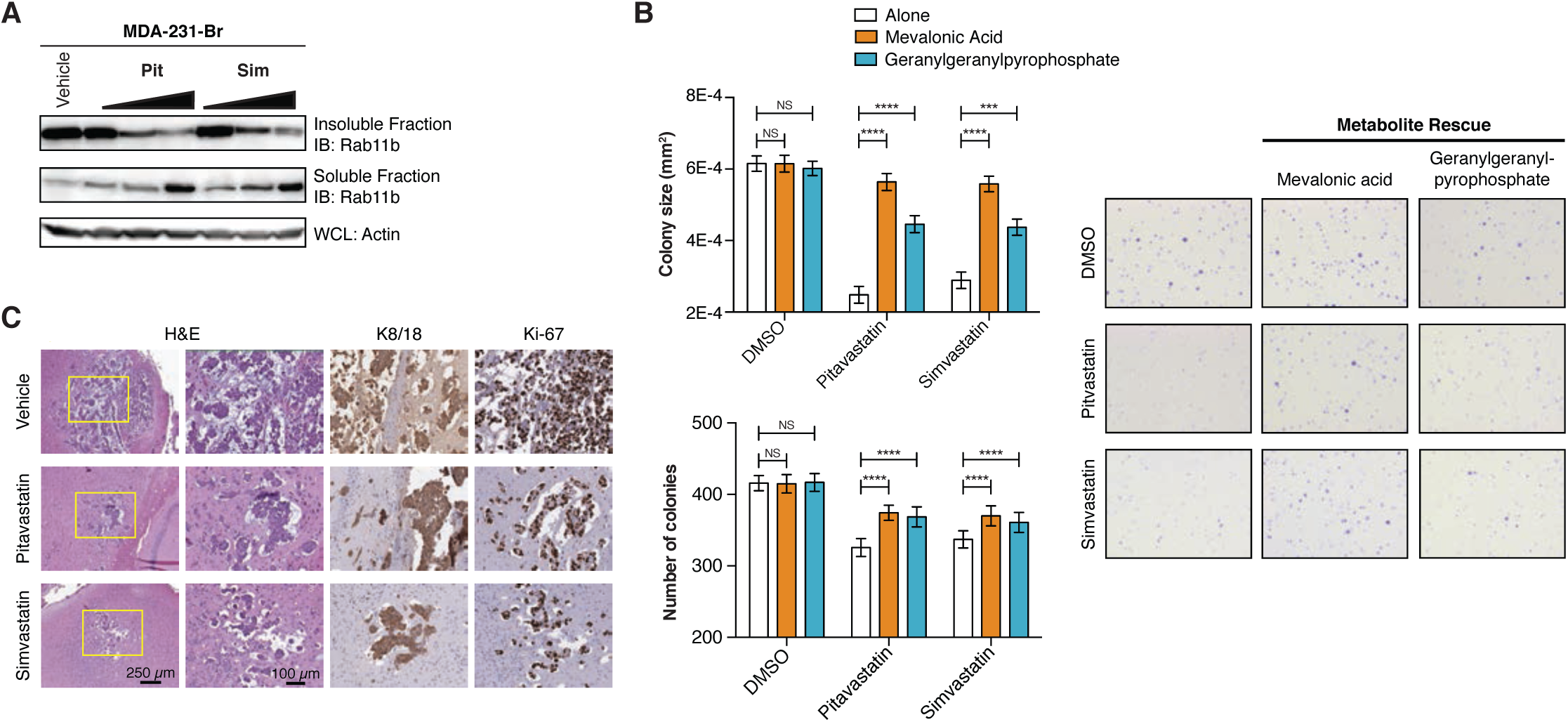
Statin inhibition of Rab11b and breast cancer brain metastasis. Related to Figure 6. **(A)** MDA-231-Br cells grown in the presence of vehicle or 10 µM-100 nM pitavastatin or simvastatin for 24 hrs. Soluble and insoluble fractions were separated with Triton X-114, and subjected to immunoblotting. **(B)** MDA-231-Br cells were grown in soft agar with 1 µM pitavastatin or simvastatin, with the addition of 100 µM mevalonic acid or 10 µM geranylgeranylpyrophosphate for two weeks. Colonies were fixed, stained and imaged. *Left*, quantification of colony number and size for ten fields per condition. Two-way ANOVA with Sidak’s multiple comparison test. *Right*, reoresentative images. **(C)** MDA-231-tdTomato cells were intracranially injected, and given daily intraperitoneal injections of vehicle or 1 mg/kg pitavastatin or 5 mg/kg simvastatin. Animals were sacrificed 34 dpi. Representative images of H&E, cytokeratin 8/18 and Ki-67 immunostaining.

## METHODS

### Cell culture

MDA-MB-231, MDA-MB-468, and BT-20 were purchased from ATCC and were maintained in DMEM/F-12 supplemented with 10% FBS and penicillin-streptomycin. MDA-MB-231-Br-EGFP was a generous gift from Patricia Steeg at the National Institute of Health (Bethesda, MD) and was maintained in DMEM/F-12 supplemented with 10% FBS and penicillin-streptomycin. BT549 and HCC38 were originally purchased from ATCC and were maintained in RPMI supplemented with 10% FBS and penicillin-streptomycin. Cancer-associated fibroblast (CAF) cell line was a generous gift from Dr. Zachary Schafer at the University of Notre Dame (Notre Dame, IN) and was maintained in RPMI supplemented with 10% FBS and penicillin-streptomycin. Primary murine glia were isolated from C57B/6 neonates using aseptic culture technique. Whole brains were passed through a 70 μm filter and spun at 500G for 5 min to remove myelin. Cells were maintained in high-glucose DMEM supplemented with 10% FBS, 10% horse serum, and penicillin-streptomycin. All cell lines were maintained at 37C with 5% CO_2_. All cells were free of mycoplasma. Human cell lines were authenticated using STR profiling by Genetica DNA Laboratories.

### Generation of cell lines

Empty vector and shRab11b constructs were transfected into HEK293T along with packaging vectors pMD2.G and psPAX2 using Lipofectamine 3000 according to manufacturer’s directions. Medium with lentiviral particles was collected 48 hr after transfection, spun down, 0.45 μm filtered, and added to recipient cell lines. 48 hr after addition of virus, cells were selected with puromycin.

### RNA-sequencing

Tumors were dissected from brain parenchyma using tdTomato fluorescence as a guide. Total RNA was isolated using the Arcturus PicoPure RNA isolation kit and sequencing libraries constructed using the Ovation RNA-Seq System, both according to manufacturer’s directions. Purified libraries were quality assessed with Qubit 2.0 Fluorometer and Agilent 2100 Bioanalyzer analysis. Paired end transcriptome sequencing (2 × 75 bp) was performed on the Illumina MiSeq sequencer at the Genomics and Bioinformatics Core Facility, University of Notre Dame. Base calling was performed using Real Time Analysis (RTA) v1.17.21.3 (Illumina). Output of RTA was converted into FastQ format with the Bcl2FastQ conversion software v1.8.4 (Illumina). Trimmed reads were aligned to reference human and mouse transcriptomes (Figure S1). Reads that aligned uniquely to the human reference genome were assigned to the tumor gene expression count table. Reads that aligned uniquely to the mouse reference genome were assigned to the microenvironment gene expression count table. All other reads were removed from downstream analysis. Differential expression was conducted using the cufflinks (Trapnell et al., 2012) and DESeq2 pipelines as described (Love et al., 2014).

### Fly stocks and genetics

The fly tester line expressing oncogenic Ras^V12^, a dsRNAi construct targeting the polarity gene **discs large)** (Dlg^RNAi^), and GFP under the control of the UAS promoter was obtained from M. Willecke (Willecke et al., 2011). The tester stock had the full genotype eyflp; UAS-Ras^V12^, UAS-Dlg^RNAi(41134)^/CyO, Gal80^(BL#9491)^; act>CD2>Gal4, UAS-GFP^S65T^ (Supplemental Figure 2B). To identify fly RNAi stocks for screening, the small, medium and large brain metastases collected at 40 dpi were treated as independent tests of the same hypothesis (that all tumors had escaped dormancy and were actively proliferating, regardless of size at the time of collection) and the combined significance was computed on a gene-wise basis using Fisher’s combined probability test. Genes with a Fisher’s combined p-value < 0.05 that were up-regulated in the 40 dpi samples were subjected to **Drosophila)** ortholog identification using the **Drosophila)** RNAi screening center (DRSC) Integrative Ortholog Prediction Tool (DIOPT) (Hu et al., 2011). For each human gene the top **Drosophila)** ortholog was identified for further analysis. Publicly available RNAi lines were then identified using the DRSC Updated Targets of RNAi Reagents tool (UP-TORR) (Hu et al., 2013). 448 RNAi lines, representing **Drosophila)** orthologs of 108 human genes were obtained from the Bloomington **Drosophila)** Stock Center and the Vienna **Drosophila)** RNAi Center for analysis.

### Fly tumor screen

For each RNAi line to be screened, 15 female virgins from the tumor tester genotype were collected and crossed to 8 males from the RNAi line. Flies were left overnight to mate and lay eggs before being passaged to a new vial. Progeny larvae from the cross were collected on the sixth day after egg laying, when larvae were wandering but not yet pupariating. Larvae were collected in 50% glycerol in water and placed at −20C for approximately 15 minutes prior to imaging. Larvae were arrayed on a plastic dish and imaged on an EVOS FL cell imaging system using the GFP filter cube and transmitted light. A minimum of 15 larvae were imaged for each cross. Transmitted light images were thresholded and used as ROIs for calculation of GFP integrated intensity. SSMD scores were calculated for each line with respect to the negative control (yw (Zhang, 2011), no RNAi), and the positive control (shPTEN (Willecke et al., 2011). Image and statistical analysis were conducted using custom MATLAB scripts.

### Animal care and use

All animal use was carried out in accordance with protocols approved by the Notre Dame Institutional Animal Care and Use Committee and were in compliance with the relevant ethical regulations regarding animal research. NOD.Cg-Rag1tm1MomIL2rgtm1Wjl/SzJ (007799/NRG) and C57BL/6J (000664/Black 6) mouse lines were purchased from The Jackson Laboratory and bred in house, with breeders refreshed directly from The Jackson Laboratory annually. For intracranial injections, 2.5 x 104 cells were injected in 690 nL Hank’s buffered saline solution, without calcium, magnesium and phenol red. Bilateral injections were made midway between bregma and lambda, 2 μm off the midline suture. For intracardiac and intracarotid injections, 2 x 105 cells were injected in 100 μL Hank’s buffered saline solution, without calcium, magnesium and phenol red. For mammary fatpad and tail vein injections, 2.5 x 105 cells were injected in 50 μL Hank’s buffered saline solution, without calcium, magnesium and phenol red. 5-10 animals were used per experimental group, and sample sizes were determined using power analysis, with expected effect size based on prior experience with metastatic animal models. For statin treatment, animals were randomly assigned to vehicle or treatment groups. Beginning two days post injection, animals were given daily intraperitoneal injections of 100 μL vehicle (0.1% hydroxypropyl methylcellulose), pitavastatin (1 mg/kg in vehicle), or simvastatin (5 mg/kg in vehicle).

### Human tissue microarray

Human tissue microarrays for normal tissue (MBN481), and brain metastases (GL861) were purchased from US Biomax.

### Immunohistochemistry

Following deparaffinization and rehydration, epitopes were retrieved by boiling slides for 10 min at 100C in sodium citrate buffer (10 mM sodium citrate, 0.05% Tween-20, pH 6.0). Slides were incubated for 1 hr at room temperature in primary antibody. Primary antibodies were detected using the VECTASTAIN Elite ABC HRP Kit (Vector Laboratories), followed by detection with ImmPACT DAB (Vector Laboratories) following manufacturer’s directions. Images were taken with an Olympus BX43.

### Immunocytochemistry

Cells were plated on glass coverslips and allowed to adhere overnight. Coverslips were fixed in 4% paraformaldehyde, permeabilized with 0.1% Triton X-100, blocked with 1% BSA, and incubated with primary antibody for 1 hr at room temperature. Primary antibodies were detected using appropriate fluorescent secondary antibodies for 1 hr at room temperature. Coverslips were stained with phalloidin for F-actin and DAPI for nuclei, and mounted. Images were taken with a Leica DM5500 for coverslips, and a Zeiss LSM 710 inverted confocal microscope for decellularized matrix.

### qPCR

Total RNA was isolated using RNAzol, and reverse transcribed using Verso cDNA Synthesis Kit. qPCR was performed using 2x SYBR Green qPCR Master Mix on an Eppendorf Mastercycler ep realplex thermocycler. Relative expression was determined using the 2CT method with logarithmic transformation.

### Rab11b activation assay

A 132 bp fragment of RAB11FIP3 corresponding to amino acids 712-756 (Junutula et al., 2004) was PCR amplified from RAB11FIP3 sequence verified cDNA and cloned into pGEX-4T-1, a generous gift from Crislyn D’Souza-Schorey, University of Notre Dame. Following sequence verification, RAB11FIP3-GST was expressed in BL21(DE)3 cells. RAB11FIP3-GST protein was purified using glutathione high-capacity magnetic beads and stored at 4C for use. For Rab11b activation assay, cell lysates were incubated with RAB11FIP3-GST beads for 1 hr at 4C, rinsed, boiled, and run on a SDS-polyacrylamide gel.

### Internalization and recycling assay

For both internalization and recycling, cells were serum starved for 30 min in medium with 0.5% BSA. Medium was replaced with 5 μg/mL transferrin-Alexa 647 in 0.5% BSA in serum-free medium. For internalization, at each timepoint cells were washed with PBS, acid stripped (0.5% glacial acetic acid, 500 μM NaCl), fixed in 4% paraformaldehyde and transferred to ice for analysis. For recycling, cells were incubated with transferrin-Alexa 647 for 1 hr at 37C, acid stripped, and medium replaced with full medium containing 50 μg/mL unlabeled transferrin. At each timepoint, cells were fixed and transferred to ice for analysis. Flow cytometry was performed on a Beckman-Coulter FC500, and analysis was performed using the FlowCore package in R FlowCore (Lo et al., 2009).

### Protein biotinylation and isolation

Surface proteins were biotinylated using the Pierce Cell Surface Protein Isolation Kit according to the manufacturer’s instructions. Briefly, surface proteins were biotinylated with Sulfo-NHS-SS-Biotin, quenched and lysed. Lysates were incubated with Neutravidin, and isolated biotinylated proteins were quantitated, boiled and run on a SDS-polyacrylamide gel. For Rab11b IP, cell surface proteins were biotinylated with Sulfo-NHS-SS-Biotin and give 24 hrs to internalize and recycle proteins. Cells were lysed using a non-denaturing lysis buffer and biotinylated proteins were pulled down with Neutravidin. Protein complexes were boiled and run on a SDS-polyacrylamide gel.

### Immunoblotting

For general immunoblotting, cells were lysed in RIPA lysis buffer and protein concentrations determined with BCA assay. Normalized lysates were boiled and run on SDS-polyacrylamide gels. To examine signaling following reinitiation of adhesion, cells were suspended for 1 hr at 37C to down-regulate adhesion-mediated signaling, and plated on cell culture plates. Cells were collected at given time points and subjected to immunoblotting as above. To examine signaling in response to poly-L-lysine and Type I Collagen, cell culture plates were coated with 0.1% poly-L-lysine or 5 μm/cm^2^ Type I Collagen for 1 hr at room temperature. Cells were suspended for 1 hr at 37C and plated on coated plates. For integrin inhibition/activation, pLKO.1 and shRab11b cells were incubated for 1 hr in suspension at 37C with 5 μg/mL P5D2 or 12G10, respectively, and plated in medium containing P5D2 or 12G10. Separation of soluble and insoluble fractions was done using Triton X-114 (Taguchi and Schätzl, 2014; Taguchi et al., 2013). Briefly, samples were lysed in 2% Triton X-114 and allowed to phase separate. The aqueous (soluble) phase was removed, and the insoluble fraction was washed several times prior to immunoblotting. All lysates were resolved with 8-15% SDS-polyacrylamide gels, and transferred to PVDF (0.2 μm, GE Healthcare) or nitrocellulose (0.45 μm, GE Healthcare) membranes as appropriate. HRP-conjugated secondaries from Cell Signaling or Pierce were used, followed by detection with SuperSignal PLUS Pico or Femto ECL.

### Soft agar assay

6 well plates were coated with 1.5 mL of 0.5% Noble agar in full medium. 25,000 cells were seeded in 0.4% Noble agar in full medium. Cells were treated as required, and medium was changed twice weekly for two or three weeks of growth. For integrin inhibition/activation, pLKO.1 and shRab11b cells were incubated for 1 hr in suspension at 37C with 5 μg/mL P5D2 or 12G10, respectively. Cultures were fed with medium containing 5 μg/mL P5D2 or 12G10 for the duration of the experiment. For statin treatment and metabolite rescue, cells were fed with medium containing 1 or 10 μM pitavastatin, simvastatin, or DMSO vehicle control for the duration of the experiment. For metabolite rescue, cells were fed with medium containing 10 μM pitavastatin or simvastatin, in addition to mevalonolactone (100 μM) or geranylgeranylpyrophosphate (10 μM) (Jiang et al., 2014) for the duration of the experiment. At endpoint, all wells were fixed with 4% paraformaldehyde, and stained with 0.05% crystal violet solution. 10 images per well were taken with a Leica MDG41 dissecting microscope.

### Proteomics - digestion and LC-MS/MS

Isolated surface proteins were run briefly on a SDS-polyacrylamide gel. LysC digestion was carried out with 1 mg of LysC for 3 h at room temperature. After adding 4 volumes of 50 mM ammonium bicarbonate buffer, 1 mg trypsin was added for tryptic digestion overnight. The next day, digestion was stopped by adding 1% TFA. Peptides were finally desalted on C18 StageTips, suspended in 10 mL 2% acetonitrile, 0.1% TFA and kept at 20C until MS analysis. The majority of samples were injected twice for MS analysis. For LC-MS analysis, a Q Exactive (Michalski et al., 2011) (Thermo Fisher Scientific) mass spectrometer was coupled on-line to an EASY-nLC 1000 HPLC system (Thermo Fisher Scientific). Desalted peptides were separated on in-house packed C18 columns (75 mm inner diameter, 50 cm length, 1.9 mm particles, Dr. Maisch GmbH, Germany) in a 250 min gradient from 2% to 60% in buffer B (80% acetonitrile, 0.5% formic acid) at 200 nL/min. Mass spectra were acquired in data-dependent mode. Briefly, each survey scan (range 300 to 1,650 m/z, resolution of 70,000 at m/z 200, maximum injection time 20 ms, ion target value of 3E6) was followed by high-energy collisional dissociation based fragmentation (HCD) of the 5 most abundant isotope patterns with a charge R 2 (normalized collision energy of 25, an isolation window of 2.2 m/z, resolution of 17,500, maximum injection time 120 ms, ion target value of 1E5). Dynamic exclusion of sequenced peptides was set to 45 s. All data was acquired using Xcalibur software (Thermo Scientific).

### Data analysis of proteomic raw files and bioinformatic analysis

MS raw files were processed with the MaxQuant software (Cox and Mann, 2008) (version 1.5.3.15). The integrated Andromeda search engine (Cox et al., 2011) was used for peptide and protein identification at an FDR of less than 1%. The human UniProtKB database (August 2015) was used as forward database and the automatically generated reverse database for the decoy search. ‘Trypsin’ was set as the enzyme specificity. We required a minimum number of 7 amino acids for the peptide identification process. Proteins that could not be discriminated by unique peptides were assigned to the same protein group (Cox and Mann, 2008) (Cox and Mann, 2008). Label-free protein quantification was performed using the MaxLFQ (Cox et al., 2014) algorithm. Briefly, quantification was based on extracted high-resolution 3D peptide features in mass-to-charge, retention time and intensity space. Only common peptides were used for pairwise ratio calculations. Protein ratios were then determined based on median peptide ratios. We required a minimum peptide ratio count of 1 to report a quantitative readout and averaged the results from duplicate measurements of the same sample. The ‘match-between-runs’ feature of MaxQuant was enabled to transfer peptide identifications across runs based on high mass accuracy and normalized retention times. Prior to data analysis, proteins, which were found as reverse hits or only identified by site modification, were filtered out. Bioinformatic analysis was performed in R (version 3.6.0) using packages available through Bioconductor (Huber et al., 2015). ENSEMBL IDs, GO terms, and GO Slim terms were annotated using biomaRt (Durinck et al., 2005, 2009). Proteins with the GO Slim “Plasma Membrane” annotation were considered surface proteins, and were used for further analysis. Pairwise t-tests with post-hoc calculation of the false discovery rate (FDR) using the Benjamini and Yekutieli procedure were conducted in R. GO term incidence was quantified, and transmembrane domains were identified using the prediction algorithm TMHMM (Krogh et al., 2001; Sonnhammer et al., 1998). Heatmaps and Cleveland plots were generated using ggplot 2 (Wickham, 2016)

### Adhesion assay

48 well plates were coated with ECM proteins for 1 hr at room temperature as follows: phenol-red free, growth factor-reduced Matrigel (5%), Type I Collagen (5 μg/cm^2^), Type IV Collagen (5 μg/cm^2^), Fibronectin (5 μg/cm^2^), Laminin (5 μg/cm^2^). Well were rinsed and cells allowed to adhere. At endpoint, cells were fixed with 10% trichloroacetic acid, and stained with 0.4% sulforhodamine blue (SRB). SRB was solubilized with 10 mM Tris, and read at 554 nm on a BioTek Synergy plate reader.

### Brain matrix decellularization and culture

Mice were anesthetized with isoflurane, and intracardially perfused with PBS. Brains were removed and cut into 1.5 mm coronal sections (Coronal Brain Matrix, Harvard Apparatus). Brains were decellularized as described (De Waele et al., 2015). Briefly, brain sections were taken through washes in 4% sodium deoxycholate, DNAse I, 3% Triton X-100, water and PBS all containing penicillin/streptomycin and amphotericin B. Removal of brain parenchymal cells was confirmed using DAPI staining as described. Decellularized brain slices were incubated in medium overnight, and cells were plated and allowed to adhere.

### Image processing and analysis

All image processing and analysis was done in ImageJ (Rueden et al., 2017) and FIJI (Schindelin et al., 2012). For integrin β_1_ activation, cell outlines were selected manually using phalloidin staining (actin) as a guide, and the integrated intensity for active integrin β_1_ (12G10) was determined. Cell area was determined in the same, using the area contained within the phalloidin boundary. Cell length was determined manually by drawing a straight line along the longest dimension of the cell. For both cell area and length, images were calibrated. To determine the number and size of soft agar colonies, calibrated images were automatically thresholded and the size and number of particles was determined. For EdU and Ki-67 quantification the number of Edu or Ki-67 positive nuclei was divided by the total number of nuclei for each field. Rab11 and integrin β_1_ IHC staining was qualitatively determined manually.

### Statistical analysis

Statistical analysis was performed using GraphPad Prism (version 6) and R (version 3.6.0). The following statistical tests were used: Fisher’s combined test, strictly standardized mean difference, two-sided t-test, ANOVA with Dunnett’s multiple comparison test, two-way ANOVA with Tukey’s multiple comparison test, analysis of contingency (Fisher’s exact test). For column and dot plots, error bars are the average +/- standard deviation unless otherwise indicated.

## Notes

**Declaration of Conflict of Interest:** The authors declare no competing interests.

